# Phylogenetic relationships in the southern African genus *Drosanthemum* (Ruschioideae, Aizoaceae)

**DOI:** 10.1101/824623

**Authors:** Sigrid Liede-Schumann, Guido W. Grimm, Nicolai M. Nürk, Alastair J. Potts, Ulrich Meve, Heidrun E.K. Hartmann

## Abstract

**Background:** *Drosanthemum,* the only genus of the tribe Drosanthemeae, is widespread over the Greater Cape Floristic Region in southern Africa. With 114 recognized species, *Drosanthemum* together with the highly succulent and species-rich tribe Ruschieae constitute the ‘core ruschioids’ in Aizoaceae. Within *Drosanthemum*, nine subgenera have been described based on flower and fruit morphology. Their phylogenetic relationships, however, have not yet been investigated, hampering understanding of monophyletic entities and patterns of geographic distribution.

**Methods:** Using chloroplast and nuclear DNA sequence data, we performed network- and tree-based phylogenetic analyses of 73 species represented by multiple accessions of *Drosanthemum*. A well-curated, geo-referenced occurrence data set comprising the phylogenetically studied and 867 further accessions was used to describe the distributional ranges of intrageneric lineages and the genus as a whole.

**Results:** Phylogenetic inference supports nine clades within *Drosanthemum*, seven of them group in two major clades, while the remaining two show ambiguous affinities. The nine clades are generally congruent to previously described subgenera within *Drosanthemum*, with exceptions such as (pseudo-) cryptic species. In-depth analyses of sequence patterns in each gene region revealed phylogenetic affinities not obvious in the phylogenetic tree. We observe a complex distribution pattern including widespread, species-rich clades expanding into arid habitats of the interior (subgenera *Drosanthemum* p.p.*, Vespertina, Xamera*) that are molecular and morphologically diverse. In contrast, less species-rich, molecularly less divergent, and morphologically unique lineages are restricted to the central Cape region and more mesic conditions (*Decidua*, *Necopina, Ossicula, Quastea, Quadrata, Speciosa*). Our results suggest initial rapid radiation generating the main lineages, with some clades showing subsequent diversification.

## Introduction

In the south-western corner of Africa, the iconic leaf-succulent Aizoaceae (ice plant family, including *Lithops*, ‘living stones’; Caryophyllales) is one of the most species-rich families in the biodiversity hot-spot of the Greater Cape Floristic Region (GCFR; Born, Linder & Desmet 2007; Mittermeier et al. 1998, 2004, 2011), ranking second in the number of endemic genera and fifth in the number of species (Manning & Goldblatt 2012). Although Aizoaceae species have received much attention both in terms of their ecology and evolution (e.g., Klak, Reeves & Hedderson 2004; Valente et al. 2014; Ellis, Weis & Gaut 2007; Hartmann 2006; Schmiedel & Jürgens 2004, Powell et al. 2019), information on phylogenetic relationships within major clades (or subfamilies) is still far from complete. Here, we aim at filling some of the knowledge-gaps providing a synoptic overview on the current classification of the family, and a study of phylogenetic relationships in the enigmatic and hitherto, phylogenetically, almost neglected genus *Drosanthemum*.

### Subfamilies of Aizoaceae: relationship of major clades

Aizoaceae currently comprises ca. 1800 species (Hartmann 2017a; Klak, Hanáček & Bruyns 2017a) classified in 145 genera and five subfamilies (Klak, Hanáček & Bruyns 2017a; Fig. 1). The first three subfamilies – Sesuvioideae, Aizooideae, Acrosanthoideae – are successive sister to Mesembryanthemoideae + Ruschioideae (Klak et al. 2003; Klak, Reeves & Hedderson 2004; Thiede 2004; Klak, Hanáček & Bruyns 2017b; for authors and species numbers see Fig. 1). Species of Mesembryanthemoideae and Ruschioideae, commonly referred to as ‘mesembs’ (Mesembryanthema; Hartmann 1991), were found in molecular phylogenetic studies to be reciprocally monophyletic (e.g., Klak et al. 2003; Thiede 2004; Klak, Hanáček & Bruyns 2017b).

**Figure 1:**
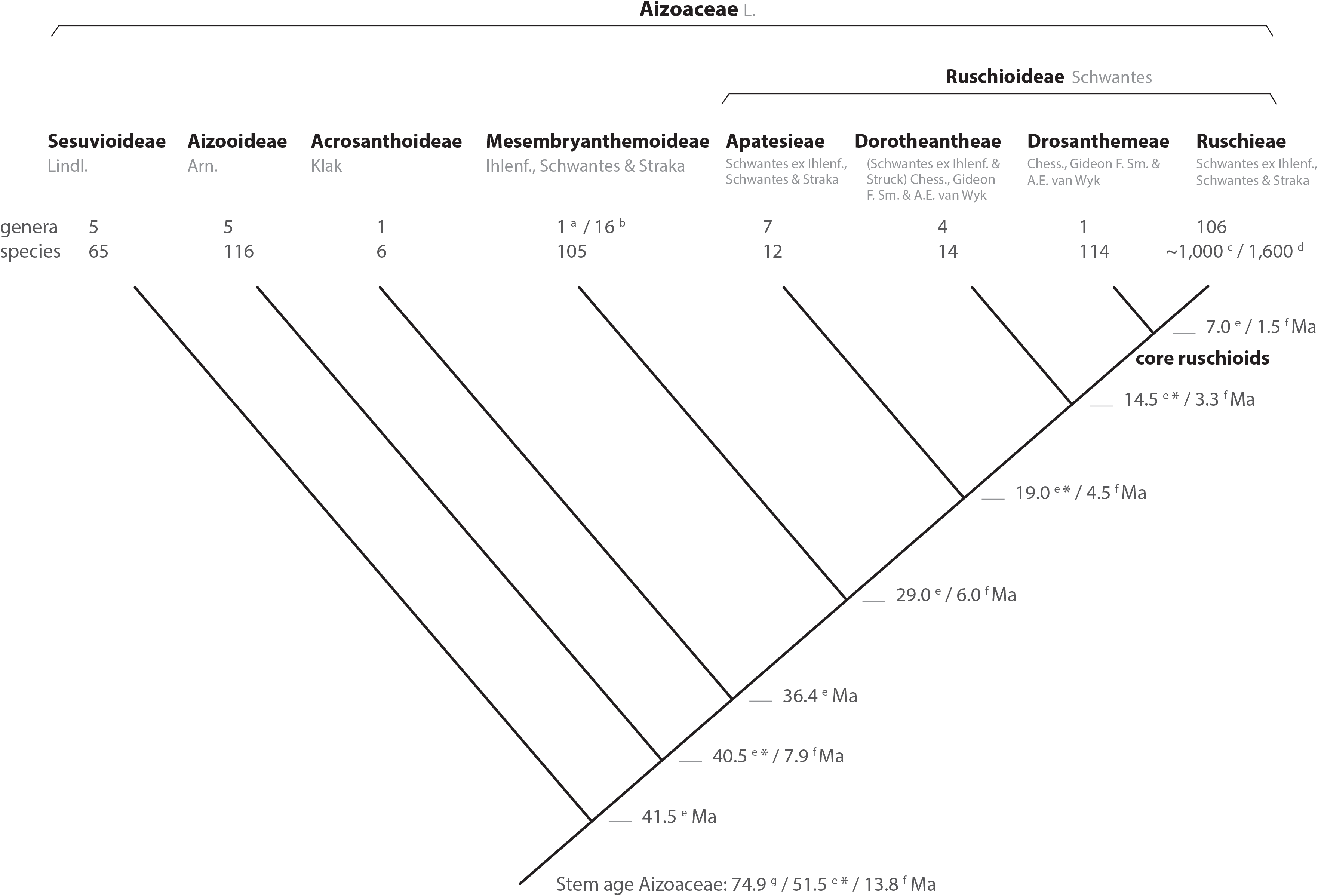
Phylogeny of Aizoaceae. A summary cladogram indicating recognized subfamilies (*sensu* Klak, Hanáček & Bruyns 2017a) and tribes (*sensu* Chesselet, Van Wyk & Smith 2004) detailing the number of genera and species and estimated node ages. Superscript letters denote reference: *a*, Klak & Bruyns 2013; *b*, Hartmann 2017a; Hartmann 2017b; *c*, Stevens, 2001 onwards; *d*, Klak, Bruyns & Hanáček 2013; *e*, Klak, Hanáček & Bruyns 2017b; *f*, Valente et al. 2014; *g*, Magallón et al. 2015. A superscript asterisk denotes ages according to Klak, Hanáček & Bruyns 2017a, Fig. S2.

Mesembryanthemoideae and Ruschioideae, as well as their sister-group relationship, are supported by morphological characters. Mesembryanthemoideae + Ruschioideae can be distinguished from the remaining Aizoaceae by raphid bundles of calcium oxalate (in contrast to calcium oxalate druses), the presence of petals of staminodial origin, half-inferior or inferior ovary and a base chromosome number of x = 9 (Bittrich & Struck 1989). The conspicuous loculicidal hygrochastic fruit capsules of ca. 98% of the species (Ihlenfeldt 1971; Parolin 2001; Parolin 2006) are lacking in Acrosanthoideae, for which xerochastic, parchment-like capsules are apomorphic, but are also predominant in subfamily Aizooideae (Bittrich 1990; Klak, Hanáček & Bruyns 2017a). In Aizooideae, however, valve wings of the capsules are either absent or very narrow, while they are well developed in Mesembryanthemoideae + Ruschioideae (Bittrich & Struck 1989).

The capsules of Mesembryanthemoideae and Ruschioideae differ in the structure of their expanding keels. The expanding keels are of purely septal origin in Mesembryanthemoideae, and mainly of valvar origin in Ruschioideae (Hartmann 1991). In floral structure, Ruschioideae are characterized almost always by crest-shaped (lophomorphic) nectaries and a parietal placentation (Hartmann & Niesler 2009), while Mesembryanthemoideae possess plain shell-shaped (coilomorphic) nectaries and a central placentation.

### Subfamilies of Aizoaceae: origin and distribution of major clades

The first-branching subfamily, Sesuvioideae (Fig. 1), originated in Africa/Arabia suggesting an African origin for the entire family (Bohley et al. 2015). While Sesuvioideae and Aizooideae dispersed as far as Australia and the Americas (Bohley et al. 2015; Klak, Hanáček & Bruyns 2017b), Acrosanthoideae, Mesembryanthemoideae and Ruschioideae are most diverse in southern Africa. Only a small number of Ruschioideae species are found outside of this area. *Delosperma* N.E.Br. is native to Madagascar and Réunion and expands with less than ten species along the East African mountains into the south-eastern part of the Arabian Peninsula (Hartmann 2016, Liede-Schumann & Newton 2018). Additionally, in Ruschioideae nine halophytic species are endemic to Australia (Prescott & Venning 1984; Hartmann 2017a; Hartmann 2017b), and possibly one species to Chile (Hartmann 2017a).

In southern Africa, most species of Acrosanthoideae, Mesembryanthemoideae and Ruschioideae are native to the Winter Rainfall Region (Verboom et al. 2009; Valente et al. 2014) in the GCFR. Acrosanthoideae with only six species is endemic to mesic fynbos, whereas Mesembryanthemoideae and Ruschioideae are speciose in more arid Succulent Karoo vegetation (Klak, Hanáček & Bruyns 2017a).

### Within Ruschioideae: relationships of major clades

Ruschioideae constitute the largest clade of Aizoaceae with estimated species richness of ca. 1600 (Stevens 2001, onwards; Klak, Bruyns & Hanáček 2013). Within Ruschioideae three tribes, Apatesieae, Dorotheantheae, and Ruschieae s.l., have been distinguished based on unique combinations of nectary and capsule characters (Chesselet, Smith & Van Wyk 2002; Fig. 1). These three tribes form well supported clades in phylogenetic analyses (Klak et al. 2003; Thiede 2004; Valente et al. 2014). Ruschieae s.l. are further characterized by the possession of wideband tracheids (Landrum 2001), endoscopic peripheral vascular bundles in the leaves (Melo-de-Pinna et al. 2014), smooth and crested mero- and holonectaries, well-developed valvar expanding tissue in the capsules (Hartmann & Niesler 2009), the loss of the *rpoC1* intron in the chloroplast DNA (cpDNA; Thiede, Schmidt & Rudolph 2007) and the possession of two *ARP* (Asymmetric Leaves1/Rough Sheath 2/Phantastica) orthologues in the nuclear DNA (Illing et al. 2009); the duplication most likely took place after the divergence of the Ruschioideae from the Mesembryanthemoideae, with the subsequent loss of one paralogue in Apatesieae and Dorotheantheae (Illing et al. 2009).

Within Ruschieae s.l. (‘core ruschioids’ *sensu* Klak, Reeves & Hedderson 2004), Klak et al. (2003) additionally revealed two clades with strong support, Ruschieae s.str. and a clade consisting only of members of *Drosanthemum* Schwantes. Species of *Delosperma*, considered closely related to *Drosanthemum* due to a papillate epidermis, often broad, flat mesophytic leaves, relatively simple hygrochastic fruits and a meronectarium have been described with *Drosanthemum* in tribe *Delospermeae* Chesselet, G. F. Smith & A. E. van Wyk (2002). In phylogenetic studies, however, *Delosperma* species group inside Ruschieae s.str. (with the exception of a few species, e.g., *Drosanthemum asperulum* and *D. longipes*, that had been shuffled between *Delosperma* and *Drosanthemum*). Consequently, Chesselet, Van Wyk & Smith (2004) included *Delosperma* in Ruschieae s.str. and coined the monogeneric Drosanthemeae as a distinct tribe sister to Ruschieae s.str. (in the following Drosanthemeae + Ruschieae = core ruschioids; Fig. 1).

While Ruschieae are characterized by fused leaf bases (Chesselet, Van Wyk & Smith 2004), an apomorphic trait is less obvious for its sister tribe Drosanthemeae. Hartmann & Bruckmann (2000) suggested capsules with a bipartite pedicel, of which the lower part appears darker due to an inner corky layer, and the upper part often thinner and agreeing in surface and colour with the capsule base. More general, species of Drosanthemeae are considered mesomorphic, compared to the highly succulent, xeromorphic Ruschieae (Klak, Bruyns & Hanáček 2013).

### Core ruschioids: relationships of lineages

A sister-group relationship of Drosanthemeae and Ruschieae has been revealed by molecular phylogenetic analyses (Klak, Bruyns & Hanáček 2013). Whether both groups are reciprocally monophyletic (and in which circumscription) is less clear (e.g., Klak, Hanáček & Bruyns 2017b). For example, molecular phylogenies identified two species erroneously included in Drosanthemeae. One of these, *Drosanthemum diversifolium* L.Bolus, was first transferred to *Knersia* H.E.K.Hartmann & Liede, a monotypic genus placed in Ruschieae (Hartmann & Liede-Schumann 2013), and later to *Drosanthemopsis* Rauschert (Ruschieae) by Klak, Hanáček & Bruyns (2018). The second species, *Drosanthemum pulverulentum* (Haw.) Schwantes, with a xeromorphic epidermis untypical for Drosanthemeae, was retrieved as member of the highly succulent clade “L1” in Ruschieae (Klak, Bruyns & Hanáček 2013; not yet formally transferred). With these corrections, Drosanthemeae comprise a single genus, *Drosanthemum*, with 114 species presently recognized and a wide distribution in the GCFR with the centre of diversity in the Cape Floristic Region (Hartmann 2017a, Van Jaarsveld 2015, 2018, Liede-Schumann, Meve & Grimm 2019).

Within *Drosanthemum*, five different floral types have been distinguished, differing mainly in number, position and relative length of petaloid staminodes (Rust, Bruckmann & Hartmann 2002, Fig. 2). Also, ten different types of capsules have been described, differing in size and shape of the capsule base and the capsule membrane, and the presence or absence of a closing body (Hartmann & Bruckmann 2000). Based on a combination of these flower and fruit types, Hartmann (2007) proposed a subdivision of *Drosanthemum* in eight subgenera. Later, Hartmann & Liede-Schumann (2014) proposed two more subgenera based on additional vegetative morphology, and also suggested the union of two of the previously described subgenera. This reflects an unusually broad variation in flower and capsule types encountered in the genus compared with other Aizoaceae genera.

**Figure 2:**
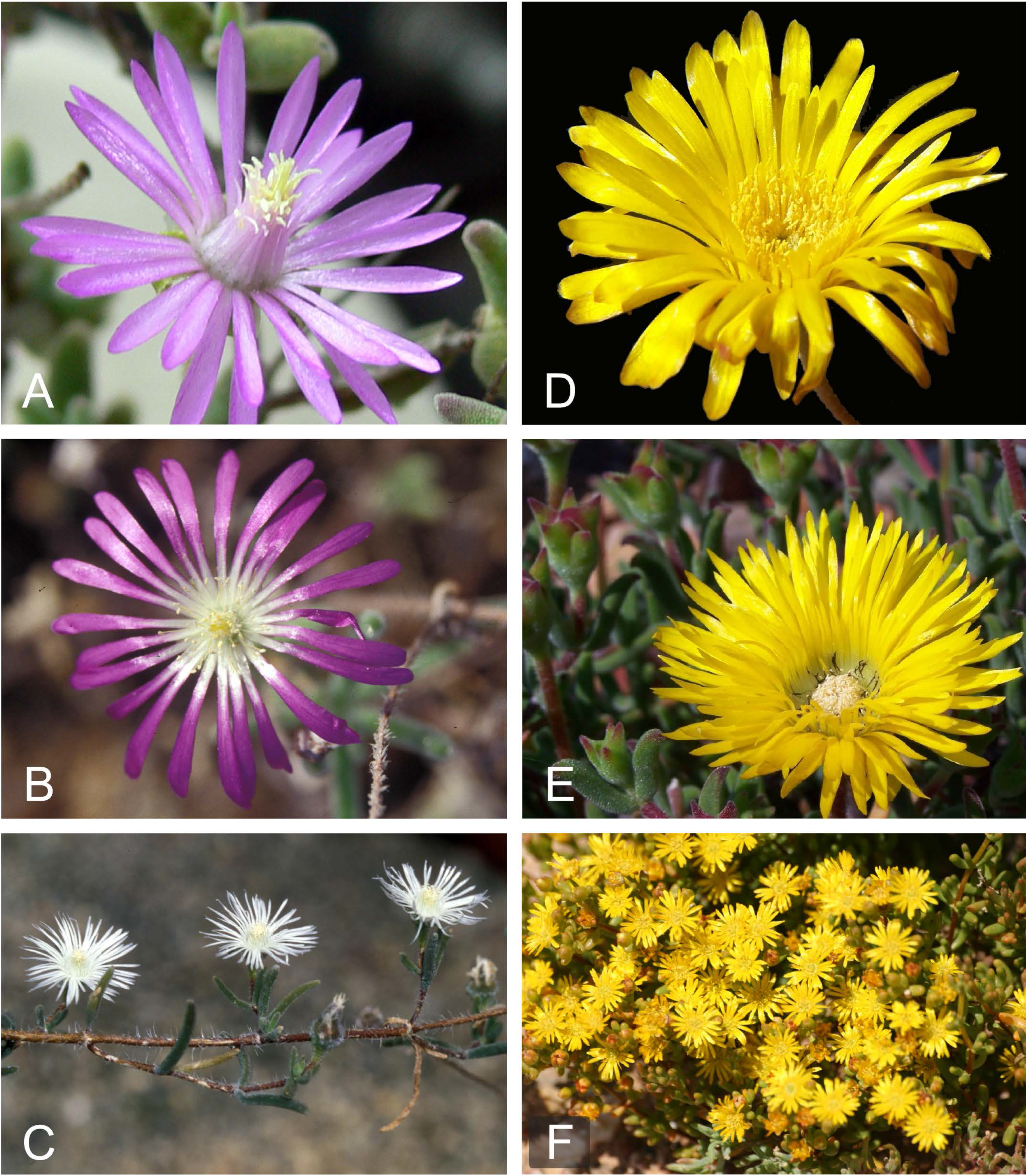
Floral diversity in *Drosanthemum*. A. *Drosanthemum lique* (subg. *Vespertina*, clade IIIa; HH 34800, HBG); B. *Drosanthemum nordenstamii* (subg. *Drosanthemum*, clade Ia; HH 31525, HBG); C. *Drosanthemum papillatum* (subg. Quastea, clade VI; HH 32425, HBG); D. *Drosanthemum cereale* (subg. *Speciosa*, clade Vb; HH 34489, HBG)—note the absence of black staminodes; E. *Drosanthemum hallii* (subg. *Speciosa*; clade Vb; HH 34610, HBG)—note the black staminodes; F. *Drosanthemum zygophylloides*. Photos A−E: H.E.K. Hartmann; F: L. Mucina.

Despite this extra-ordinary morphological diversity, molecular phylogenetic studies of *Drosanthemum* have hitherto been restricted to few species: nine species studied for ten cpDNA regions in Klak, Bruyns & Hanáček (2013) and 16 species studied for two cpDNA regions and the nuclear-encoded internal transcribed spacer (ITS) region of the 35S ribosomal DNA cistron in Hartmann & Liede-Schumann (2013). Obtaining increased species coverage representative for the phenotypic and taxonomic diversity present in *Drosanthemum* is challenging partly due to ambiguous species assignment to either *Drosanthemum* or *Delosperma* (Hartmann & Liede-Schumann 2014), but mainly due to challenges in attributing specimens to published species names in *Drosanthemum.* Ambiguous and/or overlapping diagnostic characters are common among closely related species and also present between subgenera or genera. Specimens of species flocks and (pseudo-) cryptic species (Liede-Schumann, Meve & Grimm 2019) are often hard to identify with certainty, a fact that might have hampered investigation of the genetic differentiation among *Drosanthemum* species. In this study, we build on Heidrun Hartmann’s huge field collections of identified specimens of *Drosanthemum*. The present study would not have been possible without her enduring commitment to collect, diagnose, and formally name species in the Aizoaceae.

We present a phylogenetic study of *Drosanthemum* based on a wide sampling covering more than 64% of the species richness and representing all morphology-based subgenera. We analyse chloroplast and nuclear DNA sequence variation using phylogenetic tree and network approaches and assemble a taxonomically verified *Drosanthemum* occurrence dataset covering even more of the species richness. Specifically, we ask 1) whether *Drosanthemum* constitutes a coherent, molecularly homogeneous lineage sister to Ruschiaea, or whether it forms an agglomeration (or grade) of several mesomorphic lineages; 2) whether the morphology-based subgenera represent monophyletic groups covering all species diversity in *Drosanthemum*; 3) whether the difficulties to delimit morphologically recognizable entities in *Drosanthemum* is paralleled by genetic differentiation patterns resulting in cryptic species and hitherto unrecognized relationships, or similarly, whether the most species-rich subgenus (subg. *Drosanthemum*) in fact constitutes a clade rather than a “dustbin” for species that cannot be assigned to other subgenera based on morphology; and 4) whether the clades revealed show distinct geographic distributions in the GCFR.

## Material and Methods

### Taxon sampling

We establish a collection of georeferenced *Drosanthemum* samples, which are based on sufficient material to identify key characteristics, that is, are well-identified. The full dataset (‘core collection’; n = 1000 samples) represents the most comprehensive sampling of the currently recognized 114 *Drosanthemum* species available, covering 85 species in total, with each species represented by up to 30 georeferenced samples (range, 1–30; mean, 5 samples per species). Of these we included into the molecular data set 134 accessions of *Drosanthemum*, covering 73 of the recognized species, with the more widespread and morphologically variable species represented by up to 5 accessions. To cover the full distribution in the larger subgenera, the molecular dataset comprises 21 accessions identified to subgenus, several of which most likely represent hitherto undescribed species: *Drosanthemum* (10 accessions), *Vespertina* (8 accessions), *Xamera* (2 accessions), and *Ossicula* (1 accession). Similarly, in the core collection 345 georeferenced samples identified to genus-level are included, plus 252 identified to subgenus, plus 270 identified to species. The subgeneric classification follows Hartmann (2007) and Hartmann & Liede-Schumann (2014). Geographic distribution of the subgenera corroborated with phylogenetic inference (inferred clades, see *Results*) was visualized using the RASTER library v2.8-19 (Hijmans 2019) in R v3.5.3 (R Core Team, 2019). Geographic references for the core collection are available at the Dryad digital repository (https://doi.org/10.5061/dryad.n2z34tms2; Liede-Schumann et al. 2019).

For outgroup comparison we selected a broad spectrum of species representing the three remaining tribes of Ruschioideae: Apatesieae (two accessions representing one species), Dorotheantheae (three species), and Ruschieae (49 accessions representing 47 species). We used the cpDNA dataset of Klak, Bruyns & Hanáček (2013) pruned to include one to several accessions of each Ruschieae clade (depending on clade size) with an additional nine species sequenced in previous studies of the present authors. Nuclear ITS sequences were downloaded from GenBank for accessions identical to the cpDNA data set; in five cases different accessions of the same species had to be used: *Dorotheanthus bellidiformis* (Burm.f.) N.E.Br., *Cheiridopsis excavata* L.Bolus, *Corpuscularia lehmannii* (Eckl. & Zeyh.) Schwantes, *Jacobsenia kolbei* (L.Bolus) L.Bolus & Schwantes, and *Prepodesma orpenii* (N.E.Br.) N.E.Br. A species shown by Klak, Bruyns & Hanáček (2013), to belong in Ruschieae clade L1, *Drosanthemum pulverulentum* (Haw.) Schwantes, was regarded as part of the outgroup.

### PCR and sequencing

We targeted the two cpDNA markers showing the highest intra-generic divergence between the seven *Drosanthemum* accessions used by Klak, Bruyns & Hanáček (2013), namely, the *trn*S-*trn*G intergenic spacer region and the *rpl*16 intron, using the primers and protocols provided in the original paper. In addition, the two cpDNA intergenic spacers *trnQ*–5’*rps16* and 3’*rp*S*16*–5’ *trn*K were amplified with primers trnQ^(UUG)^ and rpS16x1 and with primers rpS16x2F2 and trnK^(UUU)^, respectively (Shaw et al. 2007). The nuclear ITS region was amplified as detailed in Hassan, Thiede & Liede-Schumann (2005). Total genomic DNA was extracted from seedlings or from herbarium specimens using the DNeasy Plant MiniKit (Qiagen, Hilden, Germany), following the protocol of the manufacturer. For sequencing, the PCR products were sent to Entelechon (Regensburg, Germany) or Eurofins (Ebersberg, Germany) resulting in 473 new sequences of *Drosanthemum* species produced in this study. Forward and reverse sequences were aligned with CODONCODE ALIGNER, v.3.0.3 (CodonCode Corp., Dedham, Massachusetts, U.S.A.). Sequence data of individual marker regions were aligned with OPAL (Wheeler & Kececioglu 2007) and checked visually using MESQUITE v.3.51 (Maddison & Maddison 2018). All data used in this study is available in the Dryad digital repository https://doi.org/10.5061/dryad.n2z34tms2, Liede-Schumann et al. 2019). All sequences newly generated in this study have been submitted to ENA (for accession numbers see supplementary information S1).

### Phylogenetic analyses

Phylogenetic analyses followed our established protocols for intrageneric data sets (Khanum et al. 2016; Banag et al. 2017; Vitelli et al. 2017). We (1) used maximum likelihood (ML) tree inference and non-parametric bootstrapping (BS) analysis on (*i*) a dataset including only *Drosanthemum* species (134 accessions; ‘Drosanthemum’ dataset) to infer the ingroup topology with branch length estimates (i.e. unbiased by ingroup-outgroup branching artefacts, see next paragraph), and (*ii*) a dataset also including outgroup species (188 accessions; ‘Ruschioideae’ dataset; see *Taxon sampling*) to infer placing of *Drosanthemum* species in relation to the other Ruschioideae lineages; (2) determined the most likely *Drosanthemum* (ingroup) root using the evolutionary placement algorithm (EPA; Berger, Krompass & Stamatakis 2011) by querying a set of 47 Ruschieae species; (3) investigated competing support patterns within *Drosanthemum* using BS consensus networks (Holland & Moulton 2003; Grimm et al. 2006; Schliep et al. 2017), and (4) used median-joining (MJ; Bandelt, Forster & Röhl 1999) and statistical parsimony (SP; Templeton et al. 1992) networks to investigate within-lineage differentiation and potential molecular evolution pathways within *Drosanthemum*. Raw data and analyses code are available in the Dryad repository.

ML tree inference and BS analysis relied on RAxML v. 8.0.20 (Stamatakis 2014), set to allow for site-specific variation modelled using the ‘per-site rate’ model approximation of the Gamma distribution (Stamatakis 2006). Duplicated sequences were reduced to a single sequence resulting in 131 accessions in the ML cpDNA tree of *Drosanthemum*. To test whether *Drosanthemum* forms a grade of several lineages successively branching to Ruschieae, we conducted ML tree inference using the ‘Ruschioideae’ cpDNA data under the same RAxML settings. However, analyses of genetic differentiation among Ruschioideae species provide evidence for a fast, initial radiation within *Drosanthemum* resulting in both fast and slow-evolving lineages (indicated by branch length variation; Klak, Bruyns & Hanáček 2013). This is a data situation that may be affected by ingroup-outgroup branching artefacts (IOBA; see examples in the supplementary information S6 in Grímsson et al. 2018), a form of long-branch attraction (LBA; Bergsten 2005) potentially biasing *Drosanthemum* (ingroup) root inference.

To infer sensible outgroup-defined *Drosanthemum* roots on our data, we used outgroup-EPA implemented in RAxML following the analytical set-up of Hubert et al. (2014). EPA provides probability estimates (Berger, Krompass & Stamatakis 2011) for placing any query sequence (here: taxa representing the Ruschieae) within a given topology (here: preferred ML *Drosanthemum* cpDNA tree). Thus, it allows identifying a consensus outgroup-based root while minimising the risk of IOBAs. To identify the consensus root, we calculated a probability estimate *p*_R_ by averaging the likelihood weight ratios of query taxa per rooting scenario over all queried taxa. Several data matrices were tested in the analysis: a combined outgroup-ingroup matrix including the cpDNA used for the reference topology was run partitioned and unpartitioned, also including or excluding the most-divergent and length-polymorphic *rps*16*-trn*Q spacer region (potentially vulnerable for LBA), and including/excluding an ITS partition (the low ITS sequence variation may include signals assisting placing outgroup taxa; raw data and code are available at Dryad, Liede-Schumann et al. 2019). Because the different data matrices showed overall identical results, we present and discuss the analysis of the partitioned cpDNA dataset (all results produced are available at Dryad, Liede-Schumann et al. 2019).

BS consensus networks were computed with SplitsTree v. 4.1.13 (Huson & Bryant 2006) and was built on 1000 bootstrap (pseudo-) replicate trees generated by the ML analyses. The number of necessary BS replicates was determined using the extended majority bootstrap criterion (Pattengale et al. 2009). MJ networks for the cpDNA data were computed with NETWORK v.5.0.0.3 (Fluxus; available online www.fluxus-engineering.com/sharenet.htm) and SP networks for the ITS data with PEGAS v0.11 (Paradis 2010) in R. NETWORK was run using default settings, no character weighting was applied. Input were reduced sequence alignment matrices capturing all aspects of interspecific differentiation patterns within a clade (raw data, code, and result files of the analysis per marker are available at Dryad, Liede-Schumann et al. 2019). We differentiated four discriminating mutational patterns at the intra-clade level: (*i*) single-nucleotide polymorphisms (SNPs); (*ii*) insertions, duplications and deletions (indels), represented by a single character (gaps are treated as 5^th^ base by NETWORK by default); (*iii*) length-polymorphic sequence motifs (LP, such as multi-A motifs, which were only considered when including mutations additional to length variation); this category also includes more complex length-polymorphic patterns such as length-polymorphic AT-dominated sequence regions; and (*iv*) oligo-nucleotide motifs (ONM, short motifs with apparently linked mutations that can slightly differ in length, which were treated as a single mutational event; inversions, like the ones found in the pseudo-hairpin structure of the *trnK-rps16* spacer, are a special from of ONMs). Single-nucleotide length polymorphisms (SNLP) were not considered as they often are stochastically distributed and may be derived from sequencing or PCR errors. The input matrices were generated with PAUP* by excluding all parsimony-uninformative sites, indels, LPs, SNLPs and ONMs, and then re-including ‘key sites’ that best-represent the mutational patterns in the indels, LPs, and ONMs. In case of complex mutation patterns such as indels involving secondary SNPs or modifications, two, rarely three ‘key sites’ were exported and some cells re-coded to ensure the number of mutational events fits the situation seen in the alignment (examples shown in supplementary information S2). The highly divergent, length-polymorphic ‘high-div’ region characterising the 5’ end of the *rps*16*-trn*Q intergenic spacer, was generally excluded from the analysis but included in the haplotype documentation (Liede-Schumann et al. 2019: file Haplotyping.xlsx). The mutational patterns seen here are of high taxonomic value within clades (Liede-Schumann, Meve & Grimm 2019) but cannot be universally aligned across the entire genus, and hence, are not included for ML tree inference and BS analyses. The reasoning for using MJ and SP networks is that at the intrageneric level within subclades (‘subclade’ refers here to the nine clades within *Drosanthemum* defined in the Results section) few consistent mutations can result in a flat likelihood surface of the tree space, a situation where parsimony can be more informative than probabilistic approaches (Felsenstein 2004). In contrast to phylogenetic trees, MJ and SP haplotype networks directly depict ancestor-descendant relationships, and hence, can assist in deciding whether inferred clades in the tree are monophyletic in a strict sense, i.e. groups of inclusive common origin (Hennig 1950; see also Felsenstein 2004, chapter 10). MJ networks include all equally parsimonious solutions to a data set and produces *n*-dimensional splits graphs that can include topological alternatives. Because the MJ network used for cpDNA haplotype data can easily become diffuse or complex, especially when analysing interspecific relations, we summarized the inferred haplotypes into haplotype groups for visualizing and interpreting MJ networks (see *Results – Inter- and intra-clade differentiation patterns*).

## Results

### Genetic diversity patterns

We targeted the most variable cpDNA gene regions currently known for Aizoaceae, which provided a relatively high number of distinct alignment patterns (Table 1), although each cpDNA marker on its own provides low topological resolution (single plastid gene-region ML trees and BS consensus networks are provided in Liede-Schumann et al. 2019). Length-polymorphism was common, hence, the high proportion of gaps (undetermined cells) in the alignments, but often restricted to duplications or deletions, rarely insertions, and explicitly alignable. An exception was the *rps*16*-trn*Q intergenic spacer, which includes regions with extreme length-polymorphism and highly complex sequence patterns that are only alignable among closely related species. A notable feature is a ‘pseudo-hairpin’ sequence found in the *trn*K*-rps*16 intergenic spacer, which includes a partly clade-diagnostic strictly complementary upstream-downstream sequence pattern composed of duplications of two short sequence motifs and subsequent deletions and a “terminal” inversion (shown in the coding example in supplementary information S2, Fig. S2-A; see Liede-Schumann et al. 2019: Haplotype.xlsx, for more details).

**Table 1.** Alignment and analysis parameters for the targeted sequence regions. NLD, number of literally duplicate (identical) sequences; PUC, proportion of undetermined matrix cells (‘gappyness’); DAP, number of distinct alignment patterns; NBS, number of necessary BS pseudoreplicates; Approx. model, approximate of the DNA substitution model optimized by RAxML for each gene region (in alphabetical order: A↔C, A↔G, A↔T, C↔G, C↔T, G↔T).

| Gene region | Matrix dimension<br>(OTU × characters) | NLD | PUC | DAP | NBS | Approx.<br>model |
| --- | --- | --- | --- | --- | --- | --- |
| ITS | 112 × 440<sup>a</sup> | 27 | 2.7% | 130 | 500 | aabcde |
| *trnS-trnG* | 121 × 1157 | 34 | 30.8% | 305 | 800 | aabbba |
| *rpl16* intron | 112 × 1193 | 36 | 22.0% | 201 | 450 | aabaaa |
| *trnK-rps16* | 122 × 768 | 31 | 20.6% | 241 | 450 | aabccc |
| *rps16-trnQ* | 127 × 788<sup>b</sup> | 19 | 21.8% | 245 | 550 | aababa |

<sup>a</sup> Only ITS1 and ITS2, flanking rRNA and 5.8S rRNA genes not included. <sup>b</sup> After exclusion of the ‘high-div’ region.

In general, cpDNA sequence patterns in *Drosanthemum* are highly diagnostic at and below the level of major clades, in most cases allowing identification of haplotypes or clade-unique substitution pattern. This includes a few, potentially synapomorphic (*sensu* Hennig 1950: uniquely shared derived traits) single-base mutations in generally length-homogenous sequence portions (see Liede-Schumann et al. 2019). Indel patterns appear to be largely homoplastic, but sometimes diagnostic at the species level or for species flocks. In contrast, mutation patterns in the length-homogeneous (SNPs) and length-polymorphic regions (LP, indels, ONMs) are largely congruent with few conflicting signals for taxon splits.

The nuclear-encoded ITS region is low-divergent and shows little tree-discriminative signal, which is typical for the Aizoaceae (e.g., Klak, Bruyns & Hanáček 2013), and was not included for defining major clades and testing their coherence with the earlier proposed subgenera. Still, the genetic diversity present (Table 1) allows for the identification of more ancestral vs. more derived genotypes (supplementary information S3), which was mapped onto the cpDNA tree (Fig. 3).

**Figure 3:**
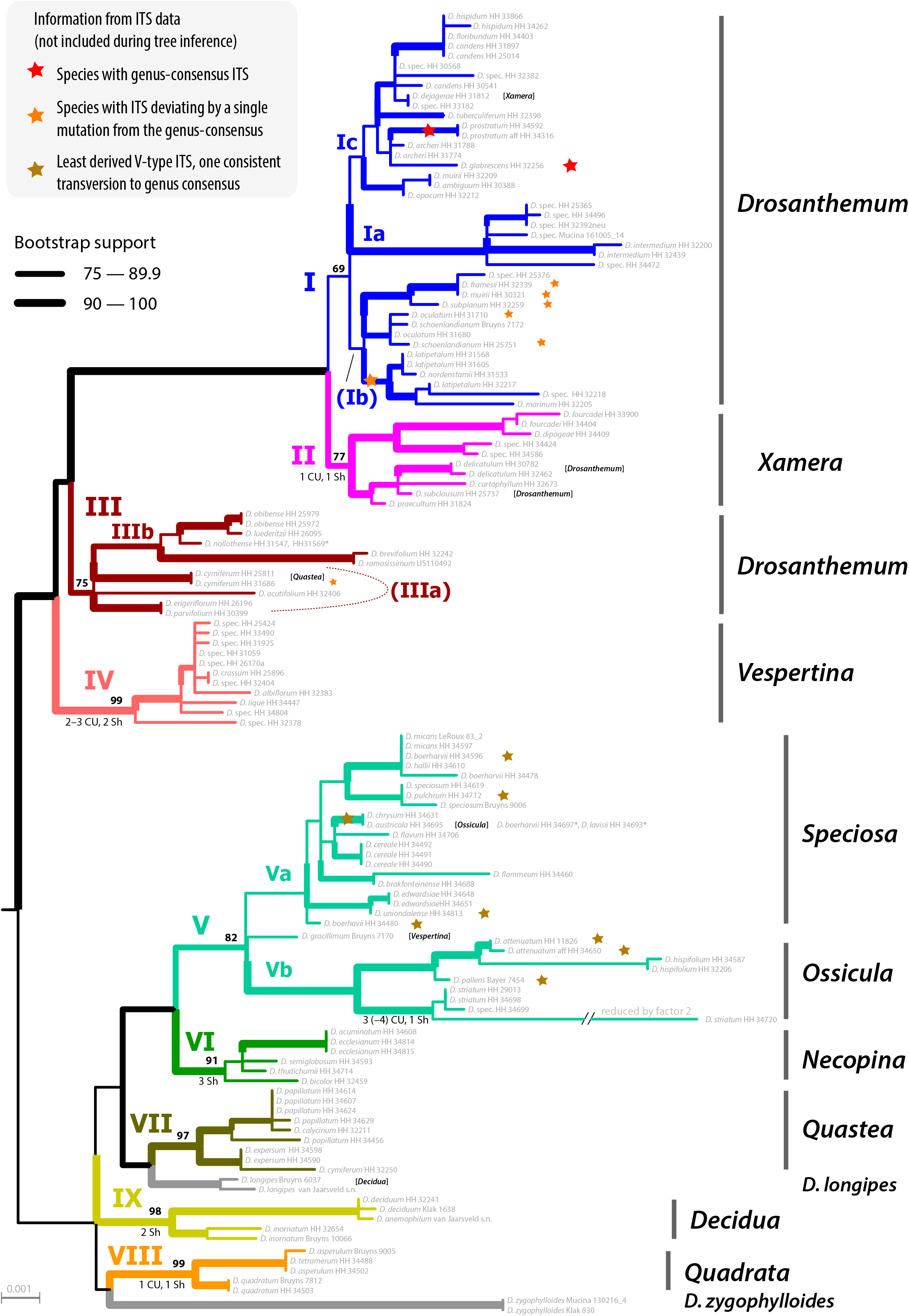
Phylogeny of *Drosanthemum*. ML tree inferred by partitioned analysis of the cpDNA sequence data. Edge lengths are scaled on expected number of substitutions per side. The nine main clades are annotated by roman numbers I–IX and coloured branches, with ML bootstrap support indicated by edge width (values given for the nine main clades). Bars and names to the right indicate subgeneric classification *sensu* Hartmann 2007. An asterisk after tip names indicate accessions with literally duplicate sequences. CU, clade unique ITS mutation pattern(s); Sh, shared ITS mutation pattern found occasionally also in other clades. Rooting is according to the most likely position (scenario 1) inferred by outgroup-EPA.

### Phylogenetic inference and potential *Drosanthemum* roots

The ML tree based on the combined cpDNA data of *Drosanthemum* indicates nine moderately (>65% BS support) to well supported clades (Fig. 3). Seven of these group in two major clades, with high support for the clade I+II+III+IV (98% BS support), addressed informally as ‘*Drosanthemum* core clade’ (Fig. 4) and low for the second clade V+VI+VII (58% BS support). Clades VIII and IX show ambiguous affinities (results not shown; for full documentation see Liede-Schumann et al. 2019). Two species, *D. longipes* (sister to clade VII) and *D. zygophylloides* (sister to VIII) are not included in the nine described clades (see *Discussion – Phylogenetic inference reflects taxonomic classification*). Notably, the nine clades overall group into six lineages (clades I–IV, V+VI, VII+*D. longipes*, VIII, XI, and *D. zygophylloides*), but without supported relationships among the six lineages. That is, earliest branching events in the ML tree are ambiguously resolved (Fig. 3)

**Figure 4:**
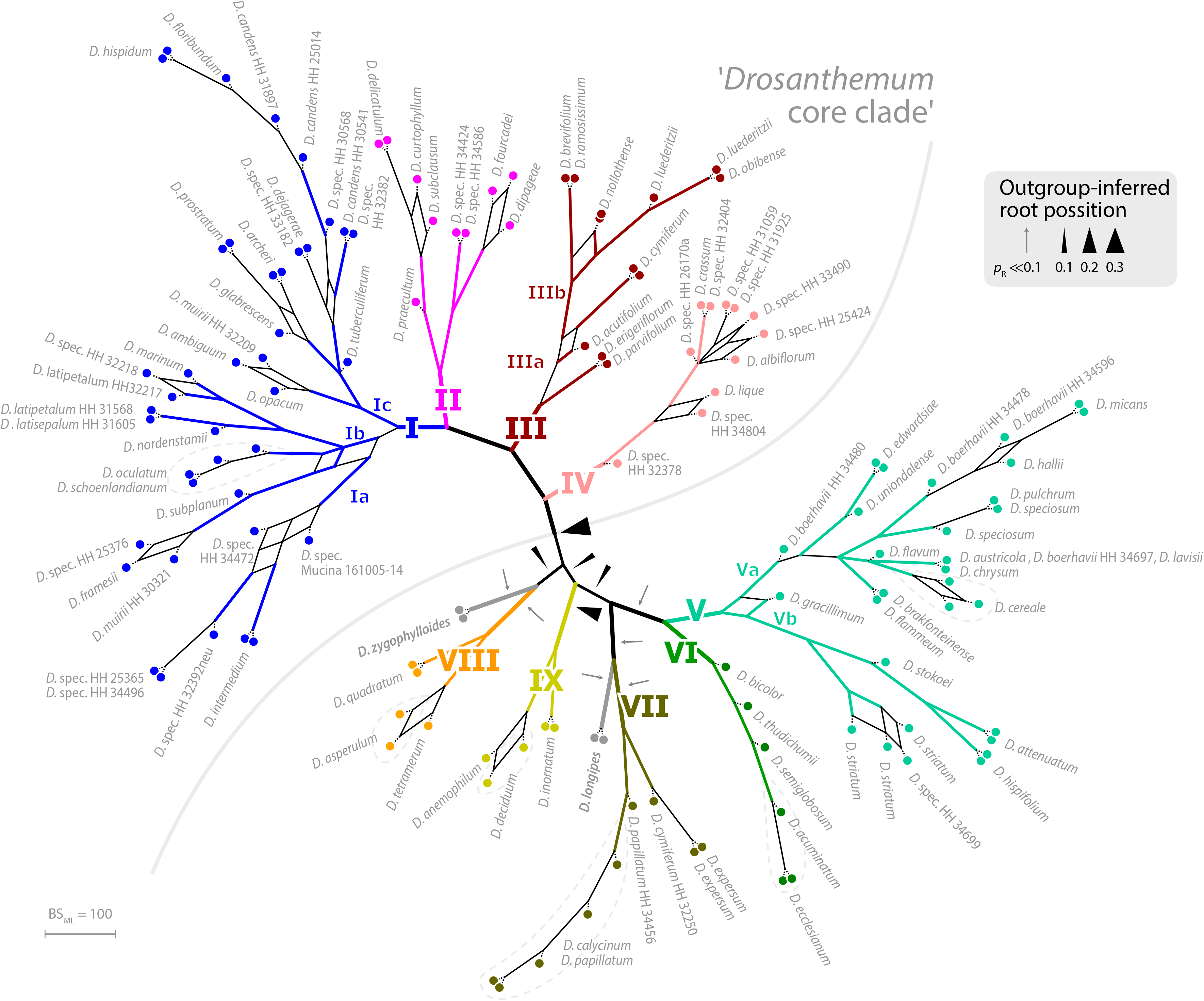
Bootstrap consensus network of *Drosanthemum*. Consensus network based on 600 pseudoreplicate samples inferred by partitioned ML analysis of the cpDNA sequence data. Edge lengths are proportional to the frequency of the phylogenetic split in the pseudoreplicate sample. Branch colours and labels are as in Fig. 3. Black arrows indicate potential root positions inferred by outgroup-EPA, with arrow size proportional to the probability estimate *p*_R_ (supplementary information S4, Table S4).

The ML tree based on the combined cpDNA data of Ruschioideae inferred monophyletic *Drosanthemum* sister to Ruschieae (100% BS support, Fig. S4.1), in accord with the ‘core ruschioids” hypothesis (Fig. 1; for details see Liede-Schumann et al. 2019). The topology within *Drosanthemum*, however, differs in parts (clade VIII and IX successive sister to the rest; not supported) from that inferred by the analysis of the combined cpDNA data of *Drosanthemum* (Fig. 3). Taken together, phylogenetic inference is consistent with a rapid initial diversification within *Drosanthemum* that was potentially too fast to leave a signal in cpDNA sequence variation in the studied markers.

Placement of the 49 queried outgroup taxa indicates eleven potential *Drosanthemum* root positions (Fig. 4), six of which are, however, unlikely considering probability estimates *p*_R_ (an order of magnitude lower), number of supporting queries (0 to 2), and phylogenetic evidence (supplementary information S4, Table S4). The remaining five root positions are summarized as follows: Scenario 1, clade I–IV sister to clade V–IX, supported by 33 queries and *p*_R_ = 0.26 (Fig. 3; S4.2); scenario 2, clade IX sister to the rest, eight queries and *p*_R_ = 0.23 (Fig. S4.3); scenario 3, clade VIII *+ D. zygophylloides* sister to the rest, one query and *p*_R_ = 0.14 (Fig. S4.4); scenario 4, cladeV–VII + *D. longipes* sister to clade I–IV + VIII + *D. zygophylloides +* IX, four queries and *p*_R_ = 0.14 (Fig. S4.5); scenario 5, clade I–IV + VIII + *D. zygophylloides* sister to clade V–IIV + *D. longipes* + IX, supported by zero queries and *p*_R_ = 0.14 (Fig. S4.6). Because the outgroup samples are notably distant in the targeted plastid gene regions to *Drosanthemum* favouring attraction of most distinct accessions, scenarios 3– 5 may be artefactual due to outgroup-ingroup (long) branch attraction. Scenario 1 is signified as the most likely root position additionally to the highest probability estimate and number of supporting queries by the distribution of ITS genotypes, indicating underived variants in clade I and V and both un- and derived ITS variants in the smaller clades outside ‘*Drosanthemum* core clade’ (Fig. 3; see also *Results, Identification of ITS genotypes*). Overall, the results obtained by outgroup-EPA are consistent with a fast radiation generating the main lineages early in the evolution of *Drosanthemum*.

### Inter- and intra-clade differentiation patterns

The haplotyping analysis of each gene region alone is in overall congruence with the combined cpDNA tree (Fig. 3). However, in some genes and/or clades coherent mutational patterns are shared by several species, which lack uniquely shared sequence patterns in other gene regions. Thus, in-detail haplotyping (Figs 5–8) further illuminates phylogenetic relationships in clades VII–IX, also including the two isolated species *D. longipes* and *D. zygophylloides*, and corroborates subgroups within clades I, III, and V (Figs. 3, 4).

**Figure 5:**
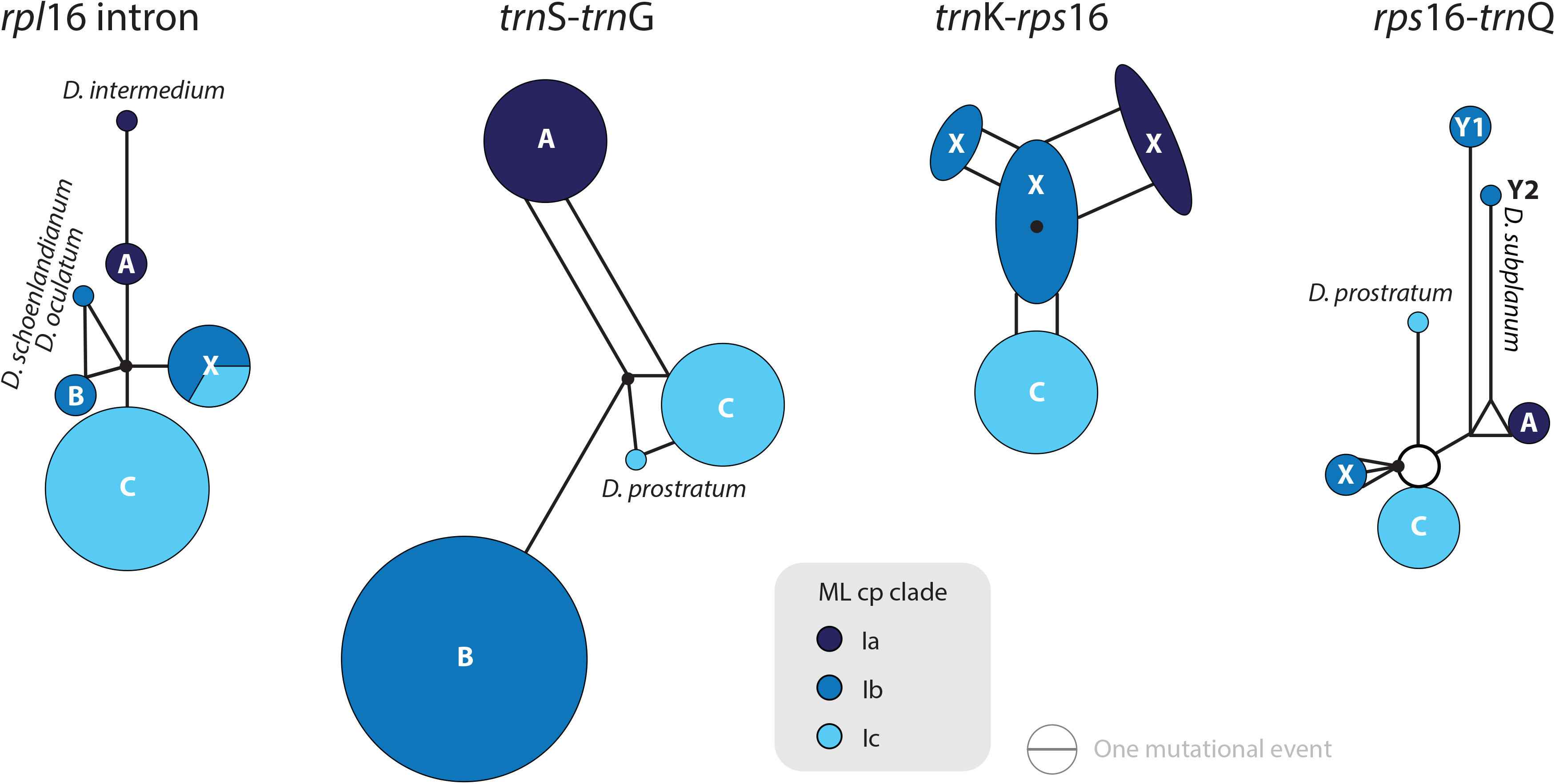
Median-joining networks of *Drosanthemum* clade I. Networks inferred by analysis of cpDNA sequence variation per chloroplast marker and simplified to haplotype groups (indicated by circles; letters in bold refer to Liede-Schumann et al. 2019: file Haplotyping.xlsx). Circle size indicate sequence divergence within the haplotype group and edge length difference in sequence variation between haplotype groups. Filled black circles denote position of the consensus sequence of the clade. Subclade Ib is paraphyletic to clades Ia and Ic according to *rpl*16 intron, *trn*K-*rps*16 and *rps*16-*trn*Q.

Clade I is divided into three subclades and the haplotyp analysis supports a monophyly of clades Ia and Ic, but not Ib (Fig. 5). Members of clade Ib are characterized by haplotypes either ancestral to those found in clades Ia and Ic (*rpl16* intron, *trnK-rps16*) or unique and strongly divergent from each other (*rps*16*-trn*Q). Clade II haplotypes are more similar to ancestral haplotypes in clade I than to those in clades III or IV. The haplotypes in clades III and IV are very similar to each other (Fig. 6). Clade III is divided into a more diverse (likely paraphyletic) grade IIIa and a monophyletic clade IIIb (Figs 3, 4, 6). Within clade III, clade IIIb forms an increasingly derived (monophyletic) lineage (*D. luederitzii + D. obibense* → *D. nollothense* → *D. brevifolium + D. ramossissimum*) that starts with grade IIIa individuals having *D. cymiferum*-like morphology but are genetically district from *D. cymiferum*. Clade V includes two sequentially coherent and mutually exclusive (reciprocally monophyletic) clades, Va and Vb (Fig. 4). In general, haplotypes of clade Vb show more uniquely shared mutational patterns than those of clade Va (Fig. 7). Figure 7 includes also the relatively similar haplotypes of the sister lineage, clade VI, which can be used to root the MJ networks (note that the edge length reflects the difference in the variable genetic patterns within clade V and does not include sequence patterns uniquely found in clade VI). Two markers, *trn*K*-rps*16 and *trn*S*-trn*G, reflect the assumed reciprocal monophyly of both clades. *Drosanthemum gracillimum* is not included in clade Va or Vb (Figs 3, 4). Only two of the considered cpDNA markers are available for this species, *trn*S*-trn*G and *rps*16*-trn*Q, with no lineage-diagnostic sequence pattern and obviously showing the putative ancestral haplotype within clade V (Fig. 7).

**Figure 6:**
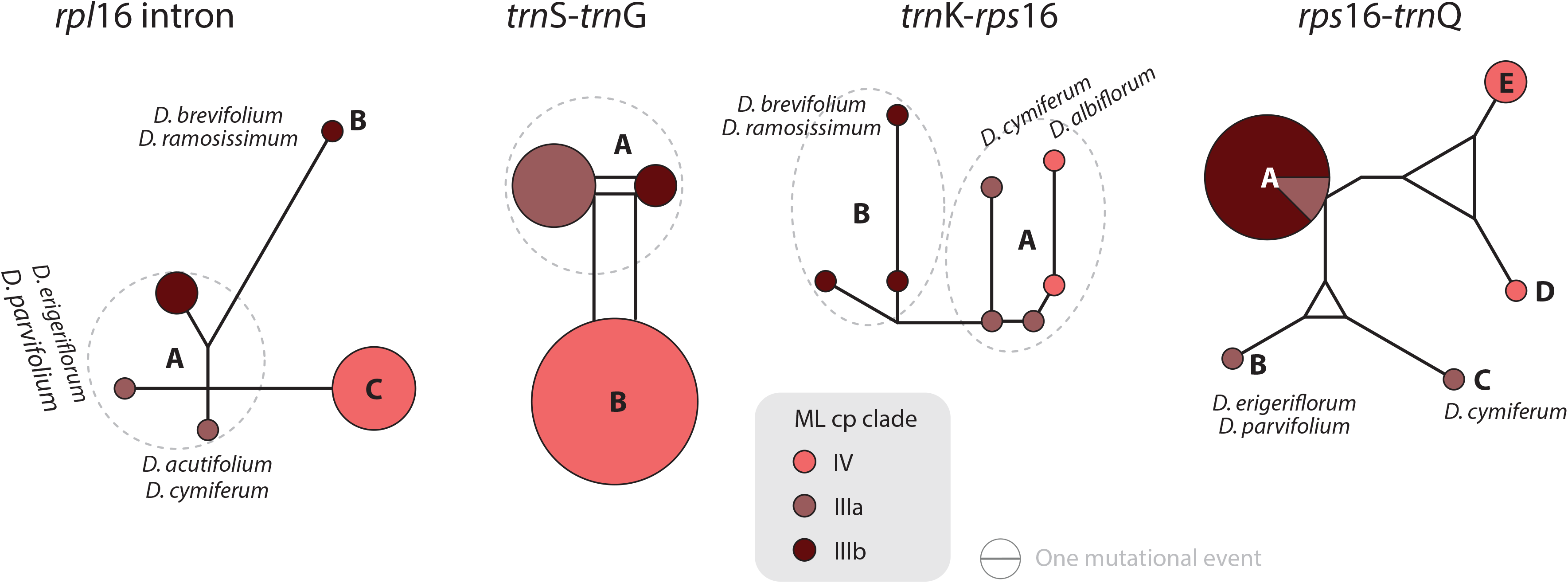
Median-joining networks of *Drosanthemum* clade III and IV. Networks inferred by analysis of cpDNA sequence variation per chloroplast marker and simplified to haplotype groups (indicated by circles; letters in bold refer to Liede-Schumann et al. 2019: file Haplotyping.xlsx). Circle size indicate sequence divergence within the haplotype group and edge length difference in sequence variation between haplotype groups. Clade IIIa bridges between haplotype groups diagnostic for clades IIIb and IV, which could be an indication of paraphyly (clade IIIa species originate from a radiation predating the formation and subsequent radiation of clades IIIb and IV).

**Figure 7:**
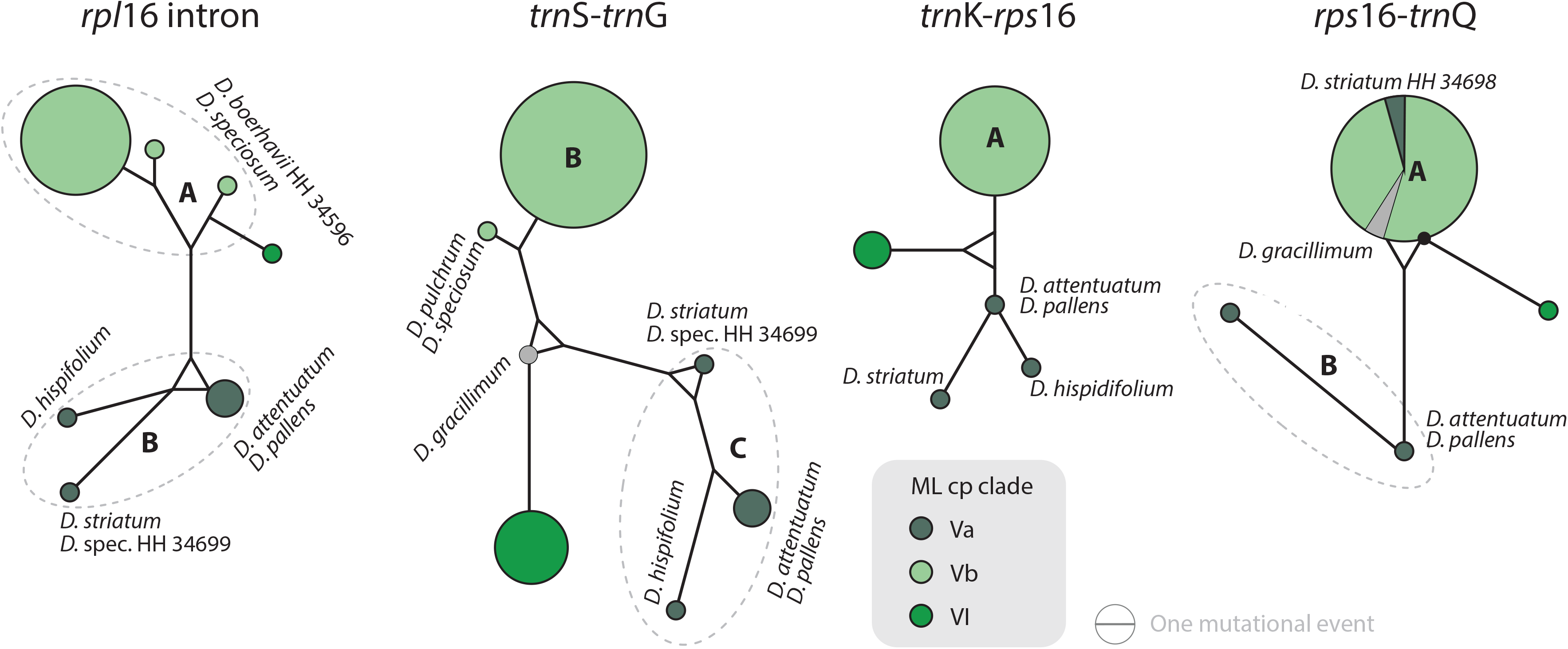
Median-joining networks of *Drosanthemum* clade V and VI. Networks inferred by analysis of cpDNA sequence variation per chloroplast marker and simplified to haplotype groups (indicated by circles; letters in bold refer to Liede-Schumann et al. 2019: file Haplotyping.xlsx). Circle size indicate sequence divergence within the haplotype group and edge length difference in sequence variation between haplotype groups. Filled black circles denote position of the consensus sequence of the clade. Note the central (*trn*S*-trn*G) or ancestral (*rps*16*-trn*Q) position of *D. gracillimum* (no *rpl*16 and *trn*K-*rps*16 data available).

Whereas haplotypes can be very divergent at the inter- and even intra-clade level (e.g. Fig. 5), they are relatively similar to each other in the smaller clades VII–IX (Fig. 8). *Drosanthemum longipes* trnS*-trn*G and *rps*16*-trn*Q haplotypes are highly similar to those of clade VII. Each gene region has a series of mutational patterns in which *D. longipes* and all members of clade VII are distinct from clade VIII and IX. In the lowest-divergent *trn*K*-rps*16 intergenic spacer region, the *D. longipes* haplotype can directly be derived from the one of clades VIII and IX (Fig. 8). *Drosanthemum longipes* is genetically closer to the putative *Drosanthemum* ancestor than to members of clade VII. In contrast, the haplotypes of *D. zygophylloides* are visibly unique within the genus (Fig. 8), which is also reflected in its long terminal branches in the cpDNA tree (Fig. 3).

**Figure 8:**
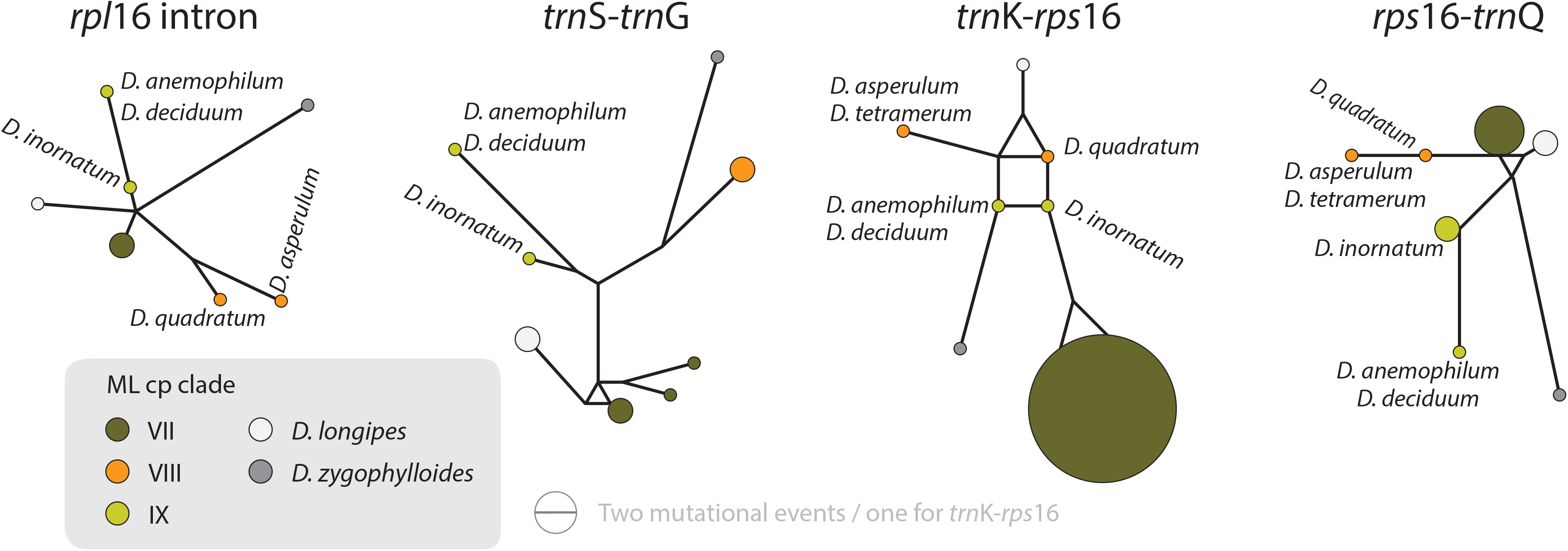
Median-joining networks of *Drosanthemum* clade VII–IX. Networks inferred by analysis of cpDNA sequence variation per chloroplast marker and simplified to haplotype groups (indicated by circles; letters in bold refer to Liede-Schumann et al. 2019: file Haplotyping.xlsx). Circle size indicate sequence divergence within the haplotype group and edge length difference in sequence variation between haplotype groups. Members of each clade are clearly differentiated but differ in the level of derivation per gene region.

### Identification of ITS genotypes in *Drosanthemum*

Analysis of nuclear ITS sequence variation reveals 62 genotypes, for which SP analysis produces an overall star-shaped network with genotypes linked to various cpDNA lineages in the centre (Fig. 9; supplementary information S3). The least derived but most common genotypes are found in distantly related clades: genotype 33 in clade I and genotype 6 clade V (Fig. 9). Genotypes 6 and 33 resemble the consensus of all ITS genotypes differing only by a single point mutation (note that genotype 33 collects several subtypes differing in an indel pattern that is ignored by the SP network; supplementary information S3; for details see Liede-Schumann et al. 2019: files Haplotype.xlsx, DataSummary.xlsx). Genotype 3 is shared by members of clade VII and IX and is central to most other (including the most common 33 and 6; Fig. 9). Clades VII–IX and the two phylogenetically isolated species, *D. longipes, D. zygophylloides*, have unique, derived genotypes. The ITS genotypes of clades II, IIIb and IV can be derived from the most ancestral ones in clade I and IIIa. The fact that ITS evolution, a stepwise derivation of putatively ancestral, consensual into clade-diagnostic, unique genotypes, can be mapped on the cpDNA phylogeny indicates that ITS differentiation in the ‘*Drosanthemum* core clade’ is in overall congruence with the cpDNA tree.

**Figure 9:**
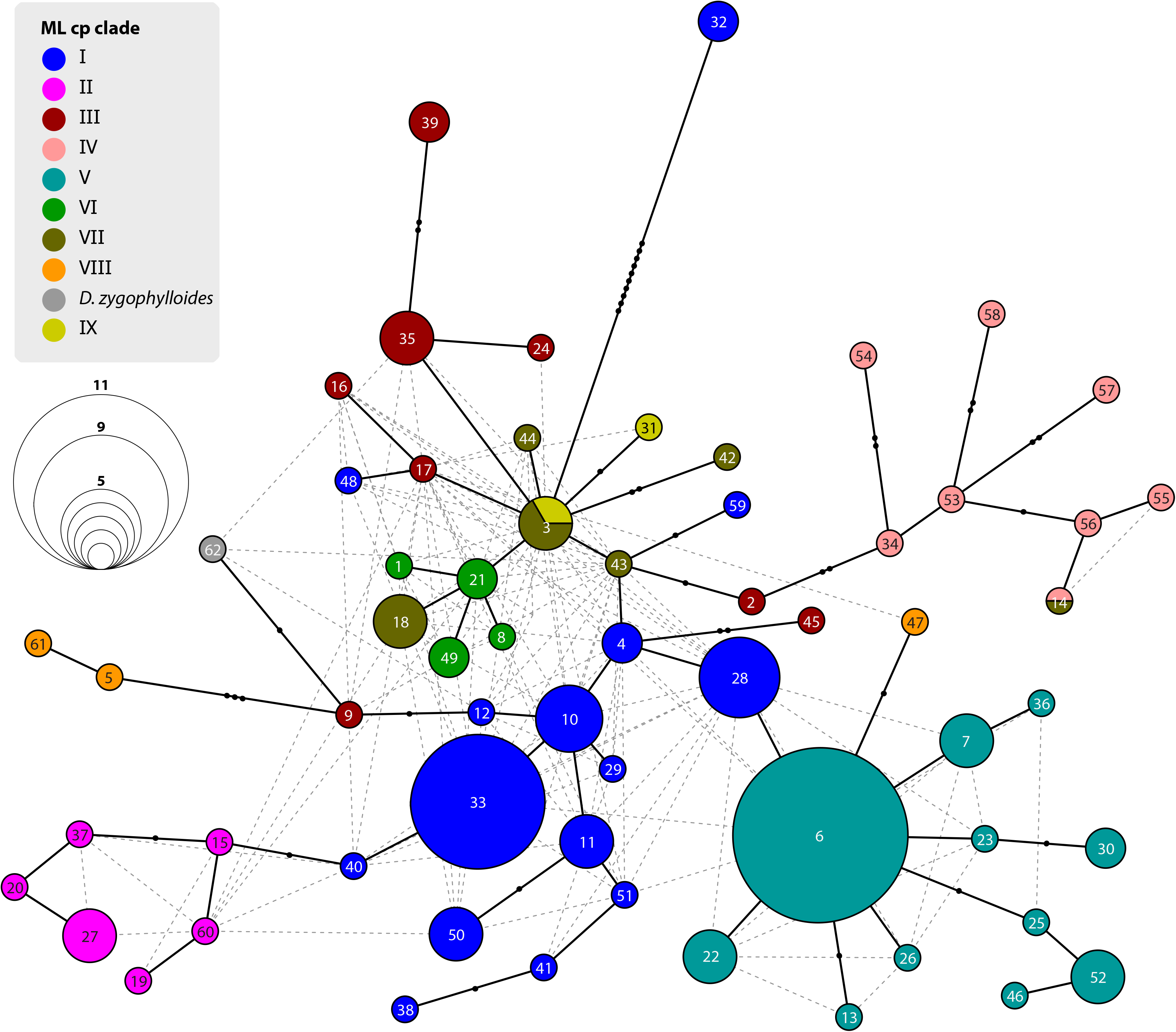
Statistical parsimony network of *Drosanthemum* ITS genotypes. Network inferred by analysis of the ITS sequence data under an infinite site model. Genotypes are indicated by circles coloured according to clades inferred by cpDNA sequence analysis (see Figs. 3, 4). Circle size indicate absolute frequency of genotypes (see legend). Black lines indicate steps in the network, filled black circles missing genotypes, and dashed grey lines alternative links. Genotypes in the centre of the graph are ancestral, those in the periphery most derived. Genotype 4 represents the genus consensus sequence found in several accessions of clade I (for details see supplementary information S3).

## Discussion

### Genetic differentiation patterns indicate fast radiation initiating diversification within *Drosanthemum*

Phylogenetic analysis, in-depth haplotyping of cpDNA, and mapping of ITS evolution on the ML cpDNA tree point towards a rapid initial diversification within the genus *Drosanthemum*. The best outgroup-EPA inferred rooting position indicates *Drosanthemum* species to group in two large clades, with clade I–IV, the ‘*Drosanthemum* core clade’, sister to clade V–IX (Fig. 3). The uncertainty in root position (Fig. 4; supplementary information S4) is consistent with a pattern expected in initial radiations (Graham & Iles 2009; Saarela et al. 2007). Nuclear ITS genetic diversity patterns indicate potential reticulation among central genotypes (polymorphic ancestral gene pool; Fig. 9, supplementary information S3), consistent with incomplete lineage sorting of ITS copies during initial radiation of *Drosanthemum* (although several other evolutionary processes in the highly repetitive nuclear ITS cistron might as well explain the pattern, e.g., incomplete concerted evolution, paralogy, intragenomic recombination, etcetera; Álvarez and Wendel, 2003; Bailey et al., 2003; Volkov et al., 2007). The star-like structure of the SP network is consistent with an initial bottleneck early in the evolution of *Drosanthemum* followed by rapid diversification (Fig. 9, supplementary information S3). Similarly, analyzed plastid sequence variation provide sufficient information to resolve nine well-supported clades within the genus *Drosanthemum*. However, the ‘backbone’ relationships among the nine clades, or to be precise, the six lineages, are not resolved (Figs. 3, 4). Taken together, the difficulties to separate and clarify the exact sequence of early branching events is a characteristic pattern in (rapid) evolutionary radiations among the plant tree of life, and has been exemplified at various phylogenetic levels, for example, in Saxifragales (Fishbein et al. 2001), within the genus *Hypericum* (Hypericaceae; Nürk et al. 2013; 2015) and in a group of South American *Lithospermum* (Boraginaceae; Weigend et al. 2010).

### Phylogenetic inference reflects taxonomic classification (as long as it does)

Within *Drosanthemum,* nine clades are revealed, which generally correspond to the recognized subgenera (Hartmann 2017a), although some exceptions exist. The deviations in morphology-based classification and phylogenetic evidence produced in this study reaveals cryptic species and several new relationships. For example, the species *D. zygophylloides*, *D. gracillimum*, and *D. longipes*, have either never been included into the subgeneric classification (*D. zygophylloides*), or phylogenetic evidence indicates affinities different from classification (*D. gracillimum*, *D. longipes*; Fig. 3). Considering our results, these species cannot be included in any of the proposed subgenera (Hartmann 2017a). Note that both *D. longipes* and the species in clade IX shed leaves in summer and resprout with the winter rains.

Subgenus *Drosanthemum* is revealed as biphyletic, with most of its species in clade I, sister to clade II (subgenus *Xamera*; see below). *Drosanthemum hispidum*, the type species of *Drosanthemum*, groups in clade I (subclade Ic; Fig. 3). The rest of the species classified in subgenus *Drosanthemum* group within clade III. No morphological diagnostic characters are obvious to distinguish the clade III species from those in clade I, and thus, clade III is not yet circumscribed as a tenth subgenus. Likewise, subgenus *Drosanthemum* species in clade I group in three subclades Ia, Ib, and Ic, but morphological characters defining these clades cannot yet be named. Hence, this species-rich subgenus is obviously biphyletic, but species assigned to it are not distributed all over the tree, i.e. subgenus *Drosanthemum* does not appear to be a “dustbin” for species that cannot be assigned based on morphology to any other subgenera.

The discussed clades I–III, together with clade IV, constitute the informally named ‘*Drosanthemum* core clade’. Clade IV corresponds to the night-flowering subgenus *Vespertina* that is characterized by flowers of the long cone type (Rust, Bruckmann & Hartmann 2002). Subgenus *Xamera* (clade II) is characterized by usually six-locular capsules and four tiny spinules below the capsule stalk and on older lateral branches (Hartmann 2007). *Drosanthemum delicatulum* and *D. subclausum* of clade II also show this character, so that their listing under subgenus *Drosanthemum* in Hartmann (2017a: 508, 532) is clearly erroneous (as is also indicated by the listing of *D. subclausum* among the species of *Xamera* in Hartmann (2017a: p 495). Conversely, *D. dejagerae* L.Bolus, attributed to *Xamera* by Hartmann (2007, 2017a) due to the presence of a six-locular capsules characteristic for the subgenus, is placed in clade Ic (subgenus *Drosanthemum* p.p.).

Of the six remaining subgenera, four, *Speciosa* (clade Va), *Ossicula* (clade Vb), *Necopina* (clade VI), and *Quastea* (clade VII), group in one clade that is, however, not well supported (Fig. 3) and also lacks obvious commonly shared, derived morphological characters. In particular, the stout and often large capsules (to 1 cm diam.) of subgenus *Speciosa* (Hartmann & Bruckmann 2000) contrast strongly with the tender and smaller capsules of the other three subgenera. However, bone-shaped closing bodies in the capsules, considered unique for subgenus *Ossicula*, have also been found in capsules of *Speciosa* species (Hartmann & Le Roux 2011), reducing their potential as a diagnostic character for *Ossicula*. This is illustrated by *D. austricola* L.Bolus, which is retrieved in subclade Va, corresponding to subgenus *Speciosa*, despite its conspicuous bone-shaped closing body, a character for which it was placed in *Ossicula* by Hartmann (2008).

While the subgeneric classification of *Drosanthemum* (Hartmann 2007; Hartmann & Liede-Schumann 2014) is largely confirmed a few unexpected placements of single species deserve mentioning. Of the three samples of *Drosanthemum cymiferum*, attributed to subgenus *Quastea* in Hartmann (2007), only one sample was retrieved in the *Quastea* clade VII, the other two in clade III (*Drosanthemum* p.p.). This species was studied in some more detail in Liede-Schumann, Meve & Grimm (2019), who did not find any consistent morphological differences between these samples and suggested a case of ‘pseudo-cryptic speciation’ (i.e. morphological analyses may find differences among these species if their potential is fully utilized; Mayr 1963; Sáez et al. 2003). A similar case is found in *D. muirii* L.Bolus, of which the two samples are retrieved with good support in subclades Ia and Ic, respectively (Fig. 3).

### Distinct geographic distributions in the Greater Cape Floristic Region

Inside the genus *Drosanthemum*, six lineages originate from a soft polytomy (precisely, they root in an unsupported part of the tree; Fig. 3), suggesting a radiation right at the start of the evolutionary history of *Drosanthemum*. To which extent this radiation was driven by ecological or geographical factors remains an open question. Interestingly, several clades comprising only 3–6 species are distributed over a restricted geographical range: clade VI *Necopina* (6 spp), clade VII *Quastea* (4 spp), and clade VIII *Quadrata* (3 spp) restricted to the western part of the Cape Mountains (Fig. 10, F–I). One species-poor lineage, clade IX *Decidua* (3 spp.), extend along the West Coast into Namibia Fig. 10, J). Species in clade V, 14 in clade Va *Speciosa* and 6 in clade Vb *Ossicula*, are almost restricted to the fynbos of GCFR (Fig. 10, F), whereas the comparatively higher species number in *Speciosa* might be the result of more thorough studies in this showy, horticulturally valuable subgenus (e.g., Hartmann 2008, Hartmann & Le Roux 2011). Notably, these clades are genetically and morphologically coherent, that is, possess unique and derived sequence patterns as well as characteristic morphologies.

**Figure 10:**
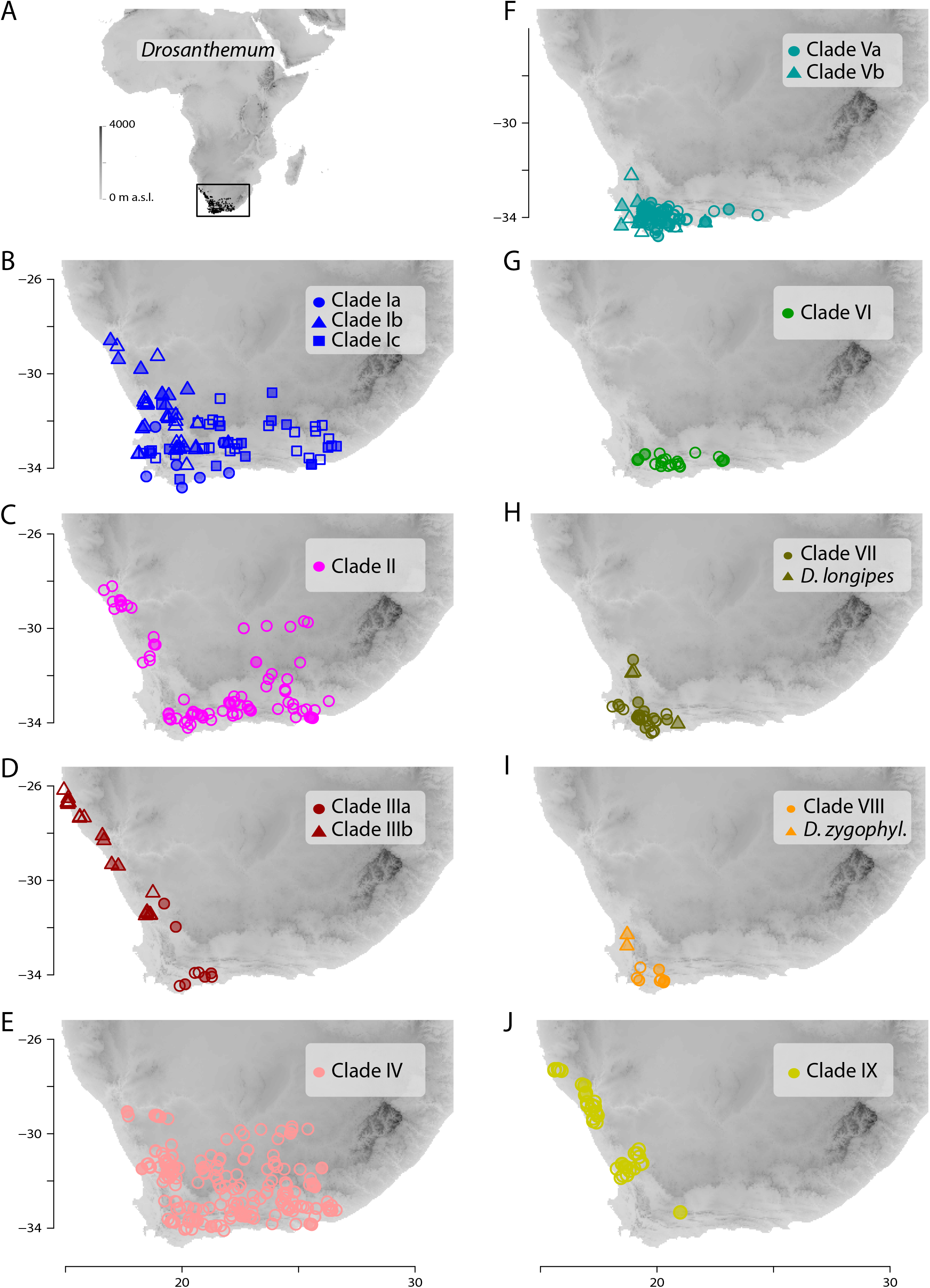
Distribution of *Drosanthemum*. A. Overall distribution of *Drosanthemum* in Africa. B–J. Clade-wise distribution of *Drosanthemum* species in southern Africa. Filled symbols indicate accessions used in the phylogeny, empty symbols indicate the remaining accessions in the occurrence dataset of *Drosanthemum*. *D. zygophyl*., *D. zygophylloides*.

The more or less narrow distribution pattern of these clades (Figs. 10, F–I) contrasts to a wide distribution of the ‘*Drosanthemum* core clade’ (Fig. 10, B– E), harbouring more widespread lineages with more species potentially indicating broader overall-habitat preferences: clade IV *Vespertina* (12 spp), clade II *Xamera* (8 spp) and the genetically and morphologically most diverse clade I (*Drosanthemum* p.p.; ≥ 55 spp; Figs. 10). The bulk of species diversity has been described in subgenus *Drosanthemum*, which falls in three subclades Ia–Ic not previously recognized (Figs. 3). These three subclades show distinct distribution patterns, with Ia restricted more or less to the fynbos area of GCFR, Ic extending far into the east and northeast, while Ib extends north to 28° S (Fig. 10, B). Clade III, composed of species hitherto considered to belong to subgenus *Drosanthemum*, shows the most diverse distribution of all clades, with a southern group of poorly resolved species, and a lineage of several species extending to the northernmost locality of *Drosanthemum*, the Brandberg in Namibia (Liede-Schumann, Meve & Grimm 2019; Fig. 10, D).

Some more species-rich clades within *Drosanthemum* have also wide ecological preferences, with representatives both at lower and higher elevations. Morphological adaptations to arid habitats are capsules with deep pockets caused by false septa enabling seed retention (Hartmann & Bruckmann 2000), which have been evolved in parallel in clade Ia and IIIb. However, whether the possession of false septa in the capsules is restricted to species of arid habitats remains an open question.

## Conclusions

In this study, we present a comprehensive phylogenetic investigation of *Drosanthemum,* a morphologically diverse genus that has so far been relatively overlooked in evolutionary studies of Aizoaceae. Our results confirm *Drosanthemum* (= Drosanthemeae) as sister lineage to Ruschieae, which is in accord with the ‘core ruschioids’ hypothesis (Klak, Reeves & Hedderson 2004; Klak, Bruyns & Hanáček 2013). Additionally, our phylogenetic evidence signifies *Drosanthemum* as a genetically well-structured but heterogenous lineage of mesomorphic plants that is, however, less species-rich than its sister clade; a pattern of diversity distribution common in the plant tree of life (Donoghue & Sanderson 2015). Still, our analysis suggest that *Drosanthemum* is not simply a depauperate lineage sister to a radiation, but instead exemplifies a radiation by itself as indicated by complex plastid and nuclear DNA sequence differentiation patterns (Figs. 3, 4, 9), and the, for Aizoaceae unusual, flower and fruit diversity present in the genus.

Occurrence patterns among the evolutionary lineages might further indicate geographic factors playing a role in species diversification in *Drosanthemum*. While most of the evolutionary history of the genus seem to have taken place in a relatively mesic environment in the southwestern parts in the GCFR, several lineages apparently have started to adapt to more arid and/or winter-cold areas. Genetically relictual species from at least two early radiations co-exist among rapidly evolving lineages, reflecting species-delimitation problems in species-rich clades. This is mirrored in in the present study that largely supports the current taxonomic concepts in *Drosanthemum* with few interesting exceptions, among others, pseudo-cryptic species.

## Supporting information

S1 Voucher Table

S2 Character recoding

S3 Statistical parsimony (SP) network analyses of ITS sequence variation

S4 Outgroup EPA

## Acknowledgements

We (SLS and HEKH) thank the Mesemb Study Group (M.S.G.) for support from the Research Fund in 2010 amd 2013. Laco Mucina (Univ. of Western Australia) is thanked for a pleasant field trip and a sample of *D. zygophylloides* and Hans-Dieter Ihlenfeldt (Univ. Hamburg) for contributing several of the outgroup samples. SLS thanks the participants of the MSc Module F1 at the University of Bayreuth from 2008 to 2015 for their work on *Drosanthemum* herbarium specimens. Angelika Täuber and Margit Gebauer (UBT) are thanked for their enduring and conscientious lab work.

## Online supporting material

**S1** [PDF]: Voucher table, indicating botanical name, voucher and ENA number of the sequences used.

**S2** [PDF]: Examples of character re-coding used for intra-clade haplotype analyses (Figures S2A–C).

**S3** [HTML]: Results of the statistical parsimony (SP) network analyses of ITS sequence variation including genotype assignment and subclade networks.

**S4** [PDF]: Results of outgroup-EPA detailing the five most probable root positions in *Drosanthemum* (Figs. S4.2–6), and the probability of all query placements (Table S4). Also including the result of the phylogenetic ML analysis of the “Ruschioideae” cpDNA data (Fig. S4.1).

