## Supplementary material for "Phylogenetic relationships in the southern African genus *Drosanthemum* (Ruschioideae, Aizoaceae)": S1 Voucher Table

**Table S1**: **Voucher**. Sequences newly produced for the present study are indicated in **bold**.

| Species | Section | Voucher | Origin | ITS | rps16-trnK | trnQ-rps16 | rpl16 | trnS-G |
|---|---|---|---|---|---|---|---|---|
| *Drosanthemum acuminatum* L.Bolus | *Necopina* | Hartmann & Bayer 34608 (HBG) | South Africa: Western Cape-Montagu | HG324008 | HG323956 | HG323982 | **LR030593** | **LR030877** |
| *Drosanthemum acutifolium* (L.Bolus) L.Bolus | *Drosanthemum* | Bruckmann & Hansen 32406 (HBG) | South Africa: Western Cape-Riversdale | **LR030506** | **LR030685** | **LR030783** | **LR030594** | **LR030878** |
| *Drosanthemum albiflorum (L.Bolus) Schwantes* | *Vespertina* | Bruckmann & Hansen 32383 (HBG) | South Africa: Western Cape-Oudtshoorn | --- | **LR030686** | **LR030784** | **LR030595** | **LR030879** |
| *Drosanthemum ambiguum* L.Bolus | *Drosanthemum* | Hartmann 30388 (HBG) | South Africa: Western Cape-Bredasdorp | --- | **LR030687** | **LR030785** | **LR030596** | **LR030880** |
| *Drosanthemum anemophilum* Van Jaarsveld & S.A.Hammer | *Decidua* | VanJaarsveld sn | South Africa: Western Cape-Laingsburg | LR030981 | LR031140 | LR031092 | LR031041 | LR031198 |
| *Drosanthemum archeri* L.Bolus | *Drosanthemum* | Hartmann et al. 31774 (HBG) | South Africa: Northern Cape-Fraserburg | **LR030507** | **LR030688** | **LR030786** | **LR030597** | **LR030881** |
| *Drosanthemum archeri* L.Bolus | *Drosanthemum* | Hartmann et al. 31788 (HBG) | South Africa: Western Cape-Prince Albert | **LR030508** | **LR030689** | **LR030787** | **LR030598** | **LR030882** |
| *Drosanthemum asperulum* (Salm-Dyck) Schwantes | *Quadrata* | Bruyns 9005 (BOL) | South Africa: Western Cape-Montagu | --- | --- | KF131854 | KF132147 | KF133114 |

| Species | Section | Voucher | Origin | ITS | rps16-trnK | trnQ-rps16 | rpl16 | trnS-G |
|---|---|---|---|---|---|---|---|---|
| *Drosanthemum asperulum* (Salm-Dyck) Schwantes | *Quadrata* | Hartmann & Bayer 34502 (HBG) | South Africa: Western Cape-Swellendam | LR030982 | LR031141 | LR031093 | LR031042 | LR031199 |
| *Drosanthemum attenuatum* (Haw.) Schwantes | *Ossicula* | Hartmann & Liede 11826 (HBG) | South Africa: Western Cape-Simonstown | LR030983 | LR031142 | LR031094 | LR031043 | LR031200 |
| *Drosanthemum attenuatum* (Haw.) Schwantes *aff.* | *Ossicula* | Hartmann & Court 34650 (HBG) | South Africa: Western Cape-Mosselbay | **LR030509** | **LR030690** | **LR030788** | **LR030599** | **LR030883** |
| *Drosanthemum austricola* L.Bolus | *Ossicula* | Hartmann & Bayer 34695 (HBG) | South Africa: Western Cape-Bredasdorp | **LR030510** | **LR030691** | **LR030789** | **LR030600** | **LR030884** |
| *Drosanthemum bicolor* L.Bolus | *Necopina* | Bruckmann & Hansen 32459 (HBG) | South Africa Western Cape-Worcester | **LR030511** | **LR030692** | **LR030790** | **LR030601** | **LR030885** |
| *Drosanthemum boerhavii* (Ecklon) H.E.K.Hartmann | *Speciosa* | Hartmann & Bayer 34697 (HBG) | South Africa: Western Cape-Swellendam | **LR030512** | **LR030693** | **LR030791** | **LR030602** | **LR030886** |
| *Drosanthemum boerhavii* (Ecklon) H.E.K.Hartmann (*D. aureopurpureum*) | *Speciosa* | Hartmann & Bayer 34478 (HBG) | South Africa: Western Cape-Swellendam | **LR030513** | **LR030694** | **LR030792** | --- | --- |
| *Drosanthemum boerhavii* (Ecklon) H.E.K.Hartmann (*D. insolitum*) | *Speciosa* | Hartmann & Bayer 34596 (HBG) | South Africa: Western Cape-Worcester | LR030984 | LR031143 | LR031095 | LR031044 | LR031201 |
| *Drosanthemum boerhavii* (*D. strictifolium* L.Bolus) | *Speciosa* | Hartmann & Bayer 34480 (HBG) | South Africa: Western Cape-Heidelberg | **LR030514** | **LR030695** | **LR030793** | **LR030603** | **LR030887** |
| *Drosanthemum brakfonteinense* Liede, A.Schweiger & H.E.K.Hartmann | *Speciosa* | Hartmann 34688 (HBG) | South Africa: Western Cape-Swellendam | **LR030515** | **LR030696** | **LR030794** | **LR030604** | **LR030888** |
| *Drosanthemum brevifolium* (Aiton) Schwantes | *Drosanthemum* | Bruckmann & Hansen 32242 (HBG) | South Africa: Western Cape-Vanrhynsdorp | **LR030516** | **LR030697** | **LR030795** | **LR030605** | **LR030889** |

| Species | Section | Voucher | Origin | ITS | rps16-trnK | trnQ-rps16 | rpl16 | trnS-G |
|---|---|---|---|---|---|---|---|---|
| *Drosanthemum calycinum* (Haw.) Schwantes | *Quastea* | Bruckmann & Hansen 32211 (HBG) | South Africa: Western Cape-Malmesbury | HG324009 | HG323957 | HG323983 | --- | **LR030890** |
| *Drosanthemum candens* (Haw.) Schwantes | *Drosanthemum* | Hartmann & Dehn 25014 (HBG) | South Africa: Eastern Cape-Albany | LR030985 | LR031144 | LR031096 | LR031045 | LR031202 |
| *Drosanthemum candens* (Haw.) Schwantes | *Drosanthemum* | Hartmann 30541 (HBG) | South Africa: Western Cape-Murraysburg | **LR030517** | **LR030698** | **LR030796** | **LR030606** | **LR030891** |
| *Drosanthemum candens* (Haw.) Schwantes | *Drosanthemum* | Hartmann et al. 31897 (HBG) | South Africa: Eastern Cape-Albany | **LR030518** | **LR030699** | **LR030797** | **LR030607** | **LR030892** |
| *Drosanthemum cereale* L.Bolus | *Speciosa* | Hartmann & Bayer 34490 (HBG) | South Africa: Western Cape-Caledon | --- | **LR030700** | **LR030798** | --- | --- |
| *Drosanthemum cereale* L.Bolus | *Speciosa* | Hartmann & Bayer 34491 (HBG) | South Africa: Western Cape-Caledon | **LR030519** | **LR030701** | --- | --- | **LR030893** |
| *Drosanthemum cereale* L.Bolus | *Speciosa* | Hartmann & Bayer 34492 (HBG) | South Africa: Western Cape-Caledon | **LR030520** | **LR030702** | **LR030799** | **LR030608** | **LR030894** |
| *Drosanthemum chrysum* L.Bolus | *Speciosa* | Hartmann & Bayer 34631 (HBG) | South Africa: Western Cape-Caledon | HG324010 | HG323958 | HG323984 | **LR030609** | **LR030895** |
| *Drosanthemum crassum* L.Bolus | *Vespertina* | Hartmann et al. 25896 (HBG) | South Africa: Western Cape-Swellendam | HG324011 | HG323959 | HG323985 | **LR030610** | **LR030896** |
| *Drosanthemum curtophyllum* L.Bolus | *Xamera* | Hartmann & Potgieter 32673 (HBG) | South Africa: Northern Cape-Namaqualand | HG324012 | HG323960 | HG323986 | **LR030611** | **LR030897** |
| *Drosanthemum cymiferum* L.Bolus | *Quastea* | Hartmann et al. 25811 (HBG) | South Africa: Northern Cape-Calvinia | **LR030521** | **LR030703** | **LR030800** | --- | **LR030898** |

| Species | Section | Voucher | Origin | ITS | rps16-trnK | trnQ-rps16 | rpl16 | trnS-G |
|---|---|---|---|---|---|---|---|---|
| *Drosanthemum cymiferum* L.Bolus | *Quastea* | Hartmann et al. 31686 (HBG) | South Africa: Northern Cape-Calvinia | **LR030522** | **LR030704** | **LR030801** | **LR030612** | **LR030899** |
| *Drosanthemum cymiferum* L.Bolus | *Quastea* | Bruckmann & Hansen 32250 (HBG) | South Africa: Northern Cape-Calvinia | **LR030523** | **LR030705** | **LR030802** | --- | **LR030900** |
| *Drosanthemum deciduum* H.E.K.Hartmann & Bruckmann | *Decidua* | Bruckmann & Hansen 32241 (HBG) | South Africa: Western Cape-Vanrhynsdorp | --- | **LR030706** | **LR030803** | **LR030613** | **LR030901** |
| *Drosanthemum deciduum* H.E.K.Hartmann & Bruckmann | *Decidua* | Klak 1638 (BOL) | South Africa: Western Cape-Vanrhynsdorp | --- | --- | KF131855 | KF132148 | KF133115 |
| *Drosanthemum dejagerae* L.Bolus | *Xamera* | Hartmann et al. 31812 (HBG) | South Africa: Western Cape-Prince Albert | **LR030524** | **LR030707** | **LR030804** | **LR030614** | **LR030902** |
| *Drosanthemum delicatulum* (L.Bolus) Schwantes | *Drosanthemum* | Hartmann 30782 (HBG) | South Africa: Western Cape-Swellendam | --- | **LR030708** | **LR030805** | **LR030615** | **LR030903** |
| *Drosanthemum delicatulum* (L.Bolus) Schwantes | *Drosanthemum* | Bruckmann & Hansen 32462 (HBG) | South Africa: Western Cape-Worcester | LR030986 | LR031145 | LR031097 | LR031046 | LR031203 |
| *Drosanthemum dipageae* H.E.K.Hartmann | *Xamera* | Hartmann 34409 (HBG) | South Africa: Eastern Cape-Uitenhage | **LR030525** | **LR030709** | **LR030806** | **LR030616** | **LR030904** |
| *Drosanthemum ecclesianum* Liede & H.E.K.Hartmann | *Necopina* | Hartmann 34814 (HBG) | South Africa: Western Cape-Uniondale | **LR030526** | **LR030710** | **LR030807** | **LR030617** | **LR030905** |
| *Drosanthemum sp. ecclesianum* Liede & H.E.K.Hartmann | *Necopina* | Hartmann 34815 (HBG) | South Africa: Western Cape-Uniondale | **LR030527** | **LR030711** | **LR030808** | **LR030618** | --- |
| *Drosanthemum edwardsiae* L.Bolus | *Speciosa* | Hartmann & Court 34648 (HBG) | South Africa: Western Cape-Mosselbay | **LR030528** | **LR030712** | **LR030809** | **LR030619** | --- |

| Species | Section | Voucher | Origin | ITS | rps16-trnK | trnQ-rps16 | rpl16 | trnS-G |
|---|---|---|---|---|---|---|---|---|
| *Drosanthemum edwardsiae* L.Bolus | *Speciosa* | Hartmann & Court 34651 (HBG) | South Africa: Western Cape-Mosselbay | HG324014 | HG323962 | HG323988 | LR031047 | --- |
| *Drosanthemum erigeriflorum* (Jacq.) Stearn | *Drosanthemum* | Hartmann & Dehn 26196 (HBG) | South Africa: Western Cape-Heidelberg | **LR030529** | **LR030713** | **LR030810** | **LR030620** | **LR030906** |
| *Drosanthemum expersum* (N.E.Br.) Schwantes | *Quastea* | Hartmann & Bayer 34590 (HBG) | South Africa: Western Cape-Ceres | **LR030530** | **LR030714** | **LR030811** | **LR030621** | **LR030907** |
| *Drosanthemum expersum* (N.E.Br.) Schwantes | *Quastea* | Hartmann & Bayer 34598 (HBG) | South Africa: Western Cape-Montagu | **LR030579** | **LR030766** | **LR030861** | **LR030667** | **LR030961** |
| *Drosanthemum flammeum* L.Bolus | *Speciosa* | Hartmann & Bayer 34460 (HBG) | South Africa: Western Cape-Worcester | HG324015 | HG323963 | HG323989 | --- | --- |
| *Drosanthemum flavum* (Haw.) Schwantes | *Speciosa* | Hartmann & Bayer 34706 (HBG) | South Africa: Western Cape-Caledon | **LR030531** | **LR030715** | **LR030812** | **LR030622** | **LR030908** |
| *Drosanthemum floribundum* (Haw.) Schwantes cf. | *Drosanthemum* | Hartmann 34403 (HBG) | South Africa: Eastern Cape-Port Elizabeth | **LR030532** | **LR030716** | **LR030813** | **LR030623** | **LR030909** |
| *Drosanthemum fourcadei* Schwantes | *Xamera* | Hartmann 33900 (HBG) | South Africa: Eastern Cape-Uitenhage | **LR030533** | **LR030717** | **LR030814** | **LR030624** | **LR030910** |
| *Drosanthemum fourcadei* Schwantes | *Xamera* | Hartmann 34404 (HBG) | South Africa: Eastern Cape-Port Elizabeth | **LR030552** | **LR030736** | **LR030832** | **LR030638** | **LR030929** |
| *Drosanthemum framesii* L.Bolus | *Drosanthemum* | Bruckmann & Hansen 32339 (UBT) | South Africa: Western Cape-Ceres | **LR030534** | **LR030718** | **LR030815** | **LR030625** | **LR030911** |
| *Drosanthemum glabrescens* L. Bolus | *Drosanthemum* | Bruckmann & Hansen 32256 (UBT) | South Africa: Northern Cape-Calvinia | **LR030535** | **LR030719** | **LR030816** | **LR030626** | **LR030912** |
| *Drosanthemum gracillimum* L.Bolus | *Vespertina* | Bruyns 7170 (BOL) | South Africa: Western Cape-Robertson | --- | --- | KF131857 | --- | KF133117 |

| Species | Section | Voucher | Origin | ITS | rps16-trnK | trnQ-rps16 | rpl16 | trnS-G |
|---|---|---|---|---|---|---|---|---|
| *Drosanthemum hallii* L.Bolus | *Speciosa* | Hartmann & Bayer 34610 (HBG) | South Africa: Western Cape-Worcester | **LR030536** | **LR030720** | **LR030817** | --- | **LR030913** |
| *Drosanthemum hispidum* (Haw.) Schwantes | *Drosanthemum* | Hartmann 33866 (HBG) | South Africa: Eastern Cape-Port Elizabeth | HG324016 | HG323964 | HG323990 | LR031048 | LR031204 |
| *Drosanthemum hispidum* (Haw.) Schwantes | *Drosanthemum* | Hartmann 34262 (HBG) | South Africa: Eastern Cape-Port Elizabeth | **LR030537** | **LR030721** | **LR030818** | **LR030627** | **LR030914** |
| *Drosanthemum hispifolium* (Haw.) Schwantes | *Ossicula* | Bruckmann & Hansen 32206 (HBG) | South Africa: Western Cape-Malmesbury | HG324017 | HG323965 | HG323991 | LR031049 | LR031205 |
| *Drosanthemum hispifolium* (Haw.) Schwantes | *Ossicula* | Hartmann & Bayer 34587 (HBG) | South Africa: Western Cape-Tulbagh | **LR030538** | **LR030722** | **LR030819** | **LR030628** | **LR030915** |
| *Drosanthemum inornatum* (L.Bolus) L.Bolus | *Decidua* | Bruyns 10066 (BOL) | Namibia: Rosh Pinah | --- | --- | KF131858 | KF132150 | KF133118 |
| *Drosanthemum inornatum* (L.Bolus) L.Bolus | *Decidua* | Hartmann & Potgieter 32654 (HBG) | South Africa: Northern Cape-Namaqualand-Numees | **LR030539** | **LR030723** | **LR030820** | --- | **LR030916** |
| *Drosanthemum intermedium* L.Bolus | | Bruckmann & Hansen 32200 (HBG) | South Africa: Western Cape-Simonstown | LR030987 | LR031146 | LR031098 | LR031050 | LR031206 |
| *Drosanthemum intermedium* L.Bolus | | Bruckmann & Hansen 32439 (HBG) | South Africa: Western Cape-Bredasdorp | **LR030540** | **LR030724** | **LR030821** | **LR030629** | **LR030917** |
| *Drosanthemum latipetalum* L.Bolus | *Drosanthemum* | Hartmann et al. 31568 (HBG) | South Africa: Northern Cape: Namaqualand | **LR030541** | **LR030725** | **LR030822** | **LR030630** | **LR030918** |
| *Drosanthemum latipetalum* L.Bolus | *Drosanthemum* | Hartmann et al. 31605 (HBG) | South Africa: Northern Cape: Namaqualand | **LR030542** | **LR030726** | **LR030823** | **LR030631** | **LR030919** |

| Species | Section | Voucher | Origin | ITS | rps16-trnK | trnQ-rps16 | rpl16 | trnS-G |
|---|---|---|---|---|---|---|---|---|
| *Drosanthemum latipetalum* L.Bolus | *Drosanthemum* | Bruckmann & Hansen 32217 (HBG) | South Africa: Western Cape-Clanwilliam | **LR030543** | **LR030727** | **LR030824** | **LR030632** | **LR030920** |
| *Drosanthemum lavisii* L.Bolus | *Speciosa* | Hartmann & Bayer 34693 (HBG) | South Africa: Western Cape-Napier | **LR030544** | **LR030728** | **LR030825** | **LR030633** | **LR030921** |
| *Drosanthemum lique* (N.E.Br.) Schwantes | *Vespertina* | Hartmann 34447 (HBG) | South Africa: Northern Cape-Williston | HG324018 | HG323966 | HG323992 | --- | **LR030922** |
| *Drosanthemum longipes* (L.Bolus) H.E.K.Hartmann | *Decidua* | Bruyns 6037 (BOL) | South Africa: Northern Cape-Calvinia | --- | --- | KF131859 | KF132151 | KF133119 |
| *Drosanthemum longipes* (L.Bolus) H.E.K.Hartmann | *Decidua* | Van Jaarsveld s.n. (HBG) | South Africa: Western Cape-Riversdale | LR030988 | LR031147 | LR031099 | LR031051 | LR031207 |
| *Drosanthemum luederitzii* (Engler) Schwantes | *Drosanthemum* | Hartmann et al. 26095 (HBG) | Namibia: Lüderitz-Süd | **LR030546** | **LR030730** | **LR030827** | **LR030635** | **LR030924** |
| *Drosanthemum marinum* L. Bolus | *Drosanthemum* | Bruckmann & Hansen 32205 (HBG) | South Africa: Western Cape-Malmesbury | **LR030547** | **LR030731** | **LR030828** | **LR030636** | **LR030925** |
| *Drosanthemum micans* (L.) Schwantes | *Speciosa* | Hartmann & Bayer 34597 (HBG) | South Africa: Western Cape-Montagu | **LR030548** | **LR030732** | **LR030829** | --- | **LR030926** |
| *Drosanthemum micans* (L.) Schwantes | *Speciosa* | LeRoux 83/2 (HBG) | South Africa: Western Cape-Worcester | **LR030549** | **LR030733** | **LR030830** | --- | **LR030927** |
| *Drosanthemum muirii* L.Bolus | *Drosanthemum* | Hartmann 30321 (HBG) | South Africa: Western Cape-Laingsburg | **LR030551** | **LR030735** | **LR030831** | **LR030637** | **LR030928** |
| *Drosanthemum muirii* L.Bolus | *Drosanthemum* | Bruckmann & Hansen 32209 (HBG) | South Africa: Northern Cape-Calvinia | **LR030550** | **LR030734** | --- | --- | --- |
| *Drosanthemum nollothense* Liede & H.E.K.Hartmann | *Drosanthemum* | Hartmann 31547 (HBG) | South Africa: Northern Cape-Namaqualand | **LR030553** | **LR030737** | **LR030833** | **LR030639** | **LR030930** |

| Species | Section | Voucher | Origin | ITS | rps16-trnK | trnQ-rps16 | rpl16 | trnS-G |
|---|---|---|---|---|---|---|---|---|
| *Drosanthemum nollothense* Liede & H.E.K.Hartmann | *Drosanthemum* | Hartmann et al. 31569 (HBG) | South Africa: Northern Cape-Namaqualand | **LR030554** | **LR030738** | **LR030834** | **LR030640** | **LR030931** |
| *Drosanthemum nordenstamii* L.Bolus | *Drosanthemum* | Hartmann et al. 31533 (HBG) | South Africa: Northern Cape-Namaqualand | **LR030555** | **LR030739** | **LR030835** | **LR030641** | **LR030932** |
| *Drosanthemum obibense* Liede & H.E.K.Hartmann | *Drosanthemum* | Hartmann et al. 25972 (HBG) | Namibia: Lüderitz-Süd-Diamanten-Sperrgebiet | **LR030545** | **LR030729** | **LR030826** | **LR030634** | **LR030923** |
| *Drosanthemum obibense* Liede & H.E.K.Hartmann | *Drosanthemum* | Hartmann et al. 25979 (HBG) | Namibia: Lüderitz-Süd | **LR030556** | **LR030740** | **LR030836** | **LR030642** | **LR030933** |
| *Drosanthemum oculatum* L.Bolus | *Drosanthemum* | Hartmann et al. 31680 (HBG) | South Africa: Northern Cape-Calvinia | **LR030557** | **LR030741** | **LR030837** | **LR030643** | **LR030934** |
| *Drosanthemum oculatum* L.Bolus | *Drosanthemum* | Hartmann et al. 31710 (HBG) | South Africa: Northern Cape-Calvinia | **LR030558** | **LR030742** | **LR030838** | --- | **LR030935** |
| *Drosanthemum opacum* L.Bolus | *Drosanthemum* | Bruckmann & Hansen 32212 (HBG) | South Africa: Western Cape-Morreesburg | **LR030559** | **LR030743** | **LR030839** | **LR030644** | **LR030936** |
| *Drosanthemum pallens* (Haw.) Schwantes (*D. stokoei*) | *Ossicula* | Bayer 7454 (HBG) | South Africa: Northern Cape: Namaqualand | HG324021 | HG323970 | HG323996 | **LR030645** | **LR030937** |
| *Drosanthemum papillatum* L.Bolus | *Quastea* | Hartmann 34456 (HBG) - Worcester | South Africa: Western Cape-Worcester | **LR030560** | **LR030744** | **LR030840** | **LR030646** | **LR030938** |
| *Drosanthemum papillatum* L.Bolus | *Quastea* | Hartmann & Bayer 34607 (HBG) | South Africa: Western Cape-Swellendam | HG324020 | HG323968 | HG323994 | LR031052 | LR031208 |
| *Drosanthemum papillatum* L.Bolus | *Quastea* | Hartmann & Bayer 34614 (HBG) | South Africa: Western Cape-Worcester | **LR030561** | **LR030745** | **LR030841** | **LR030647** | **LR030939** |

| **Species** | **Section** | **Voucher** | **Origin** | **ITS** | **rps16-trnK** | **trnQ-rps16** | **rpl16** | **trnS-G** |
|---|---|---|---|---|---|---|---|---|
| *Drosanthemum papillatum* L.Bolus | *Quastea* | Hartmann & Bayer 34624 (HBG) | South Africa: Western Cape-Montagu | **LR030562** | **LR030746** | **LR030842** | **LR030648** | **LR030940** |
| *Drosanthemum papillatum* L.Bolus | *Quastea* | Hartmann & Bayer 34629 (HBG) | South Africa: Western Cape-Caledon | --- | **LR030747** | **LR030843** | --- | **LR030941** |
| *Drosanthemum parvifolium* (Haw.) Schwantes | *Drosanthemum* | Hartmann 30399 (HBG) | South Africa: Western Cape-Bredasdorp | **LR030563** | **LR030748** | **LR030844** | **LR030649** | **LR030942** |
| *Drosanthemum praecultum* (N.E.Br. ) Schwantes (*D. montaguense*) | *Xamera* | Hartmann et al. 31824 (HBG) | South Africa: Western Cape-Uniondale | HG324019 | HG323967 | HG323993 | **LR030650** | **LR030943** |
| *Drosanthemum prostratum* L.Bolus *aff.* | *Drosanthemum* | Hartmann 34316 (HBG) | South Africa: Northern Cape-Calvinia | **LR030564** | **LR030749** | **LR030845** | **LR030651** | **LR030944** |
| *Drosanthemum prostratum* L.Bolus | *Drosanthemum* | Hartmann & Bayer 34592 (HBG) | South Africa: Western Cape-Ceres | **LR030565** | **LR030750** | **LR030846** | **LR030652** | **LR030945** |
| *Drosanthemum pulchrum* L.Bolus | *Speciosa* | Hartmann & Bayer 34712 (HBG) | South Africa: Western Cape-Worcester | **LR030566** | **LR030751** | **LR030847** | --- | **LR030946** |
| *Drosanthemum quadratum* Klak | *Quadrata* | Bruyns 7812 (BOL) | South Africa: Western Cape-Swellendam | --- | --- | KF131861 | KF132153 | KF133121 |
| *Drosanthemum quadratum* Klak | *Quadrata* | Hartmann & Bayer 34503 (HBG) | South Africa: Western Cape-Swellendam | **LR030568** | **LR030753** | **LR030849** | **LR030654** | **LR030948** |
| *Drosanthemum ramosissimum* L.Bolus | *Drosanthemum* | Schmiedel 110492 (HBG) | South Africa: Western Cape-Moedverloren | **LR030569** | **LR030754** | **LR030850** | **LR030655** | **LR030949** |
| *Drosanthemum schoenlandianum* L.Bolus | *Drosanthemum* | Bruyns 7172 (BOL) | | AJ438214 | --- | JN896435 | KF132154 | JN896382 |
| *Drosanthemum schoenlandianum* L.Bolus | *Drosanthemum* | Hartmann et al. 25751 (HBG) | South Africa: Northern Cape-Calvinia | LR030989 | LR031148 | LR031100 | LR031053 | --- |

| Species | Section | Voucher | Origin | ITS | rps16-trnK | trnQ-rps16 | rpl16 | trnS-G |
|---|---|---|---|---|---|---|---|---|
| *Drosanthemum semiglobosum* L.Bolus | *Necopina* | Hartmann & Bayer 34593 (HBG) | South Africa: Western Cape-Worcester | **LR030570** | **LR030755** | **LR030851** | **LR030656** | **LR030950** |
| *Drosanthemum sp.* | *Drosanthemum* | Bruckmann & Hansen 32218 (HBG) | South Africa: Western Cape-Clanwilliam | **LR030571** | **LR030756** | **LR030852** | **LR030657** | **LR030951** |
| *Drosanthemum sp.* | *Drosanthemum* | Bruckmann & Hansen 32382 (HBG) | South Africa: Western Cape-Oudtshoorn | --- | **LR030757** | **LR030853** | **LR030658** | **LR030952** |
| *Drosanthemum sp.* | *Drosanthemum* | Bruckmann & Hansen 32392 (HBG) | South Africa: Western Cape-Mosselbay | **LR030572** | **LR030758** | **LR030854** | **LR030659** | **LR030953** |
| *Drosanthemum sp.* | *Drosanthemum* | Hartmann & Bayer 34472 (HBG) | South Africa: Western Cape-Robertson | **LR030573** | **LR030759** | **LR030855** | **LR030660** | **LR030954** |
| *Drosanthemum sp.* | *Drosanthemum* | Hartmann & Bayer 34496 (HBG) | South Africa: Western Cape-Swellendam | **LR030574** | **LR030760** | **LR030856** | **LR030661** | **LR030955** |
| *Drosanthemum sp.* | *Drosanthemum* | Hartmann 30568 (HBG) | South Africa: Eastern Cape-Graaff-Reinet | **LR030575** | **LR030761** | **LR030857** | **LR030662** | **LR030956** |
| *Drosanthemum sp.* | *Drosanthemum* | Hartmann 33182 (HBG) | South Africa: Northern Cape-De Aar | **LR030576** | **LR030762** | **LR030858** | **LR030663** | **LR030957** |
| *Drosanthemum sp.* | *Drosanthemum* | Hartmann et al. 25365 (HBG) | South Africa: Western Cape-Prince Albert | **LR030577** | **LR030763** | **LR030859** | **LR030664** | **LR030958** |
| *Drosanthemum sp.* | *Drosanthemum* | Hartmann et al. 25376 (HBG) | South Africa: Western Cape-Ceres | --- | **LR030764** | **LR030860** | **LR030665** | **LR030959** |
| *Drosanthemum sp.* | *Ossicula* | Hartmann & Bayer 34699 (HBG) | South Africa: Western Cape-Swellendam | **LR030578** | **LR030765** | **LR215984** | **LR030666** | **LR030960** |

| Species | Section | Voucher | Origin | ITS | rps16-trnK | trnQ-rps16 | rpl16 | trnS-G |
|---|---|---|---|---|---|---|---|---|
| *Drosanthemum sp.* | *Vespertina* | Bruckmann & Hansen 32378 (HBG) | South Africa: Western Cape-Prince Albert | **LR030580** | **LR030767** | **LR030862** | **LR030668** | **LR030962** |
| *Drosanthemum sp.* | *Vespertina* | Bruckmann & Hansen 32404 (HBG) | South Africa: Western Cape-Riversdale | **LR030581** | **LR030768** | **LR030863** | **LR030669** | **LR030963** |
| *Drosanthemum sp.* | *Vespertina* | Hartmann 31059 (HBG) | South Africa: Northern Cape-Philipstown | **LR030582** | **LR030769** | **LR030864** | **LR030670** | **LR030964** |
| *Drosanthemum sp.* | *Vespertina* | Hartmann 34804 (HBG) | South Africa: Eastern Cape-Port Elizabeth | **LR030583** | **LR030770** | **LR030865** | **LR030671** | **LR030965** |
| *Drosanthemum sp.* | *Vespertina* | Hartmann et al. 25424 (HBG) | South Africa: Northern Cape-Namaqualand | --- | **LR030771** | **LR030866** | **LR030672** | **LR030966** |
| *Drosanthemum sp.* | *Vespertina* | Hartmann et al. 26170a (HBG) | South Africa: Western Cape-Vredendal | **LR030584** | **LR030772** | **LR030867** | **LR030673** | **LR030967** |
| *Drosanthemum sp.* | *Vespertina* | Hartmann et al. 31925 (HBG) | South Africa: Eastern Cape-Cradock | --- | **LR030773** | **LR030868** | **LR030674** | **LR030968** |
| *Drosanthemum sp.* | *Vespertina* | Hartmann et al. 33490 (HBG) | South Africa: Eastern Cape-Hofmeyr | **LR030585** | **LR030774** | **LR030869** | **LR030675** | **LR030969** |
| *Drosanthemum sp.* | *Xamera* | Hartmann & Milton 34586 (HBG) | South Africa: Western Cape-Prince Albert | **LR030586** | **LR030775** | **LR030870** | **LR030676** | **LR030970** |
| *Drosanthemum sp.* | *Xamera* | Hartmann 34424 (HBG) | South Africa: Northern Cape-Victoria West | **LR030587** | **LR030776** | **LR030871** | **LR030677** | **LR030971** |
| *Drosanthemum sp.* | *Drosanthemum* | Mucina 161005-14 (NBG) | South Africa: Western Cape-Piketberg | **LR030588** | **LR030777** | **LR030872** | **LR030678** | **LR030972** |

| Species | Section | Voucher | Origin | ITS | rps16-trnK | trnQ-rps16 | rpl16 | trnS-G |
|---|---|---|---|---|---|---|---|---|
| *Drosanthemum speciosum* (Haw.) Schwantes | *Speciosa* | Bruyns 9006 (BOL) | South Africa: Western Cape-Worcester | --- | --- | KF131862 | KF132155 | KF133122 |
| *Drosanthemum speciosum* (Haw.) Schwantes | *Speciosa* | Hartmann & Bayer 34619 (HBG) | South Africa: Western Cape-Worcester | HE585039 | HG323969 | HG323995 | --- | --- |
| *Drosanthemum striatum* Schwantes | *Ossicula* | Hartmann 29013 (HBG) | South Africa: Western Cape-Worcester | **LR030589** | **LR030778** | --- | **LR030679** | **LR030973** |
| *Drosanthemum striatum* Schwantes | *Ossicula* | Hartmann & Bayer 34698 (HBG) | South Africa: Western Cape-Swellendam | **LR030567** | **LR030752** | **LR030848** | **LR030653** | **LR030947** |
| *Drosanthemum striatum* Schwantes | *Ossicula* | Hartmann & Bayer 34720 (HBG) | South Africa: Western Cape-Worcester | **LR030590** | **LR030779** | **LR030873** | **LR030680** | **LR030974** |
| *Drosanthemum subclausum* L.Bolus | *Drosanthemum* | Hartmann et al. 25737 (HBG) | South Africa: Western Cape-Vanrhynsdorp | HG324022 | HG323971 | HG323997 | **LR030681** | **LR030975** |
| *Drosanthemum subplanum L.Bolus* | *Drosanthemum* | Bruckmann & Hansen 32259 (HBG) | South Africa: Northern Cape-Calvinia | **LR030591** | **LR030780** | **LR030874** | **LR030682** | **LR030976** |
| *Drosanthemum tetramerum* H.E.K.Hartmann | *Quadrata* | Hartmann & Bayer 34488 (HBG) | South Africa: Western Cape-Caledon | LR030990 | LR031149 | LR031101 | --- | --- |
| *Drosanthemum thudichumii* L.Bolus | *Necopina* | Hartmann & Bayer 34714 (HBG) | South Africa: Western Cape-Worcester | HG324023 | HG323972 | HG323998 | LR031054 | LR031209 |
| *Drosanthemum tuberculiferum* L. Bolus | *Drosanthemum* | Bruckmann & Hansen 32398 (HBG) | South Africa: Western Cape-Riversdale | --- | **LR030781** | **LR030875** | **LR030683** | **LR030977** |
| *Drosanthemum uniondalense* H.E.K.Hartmann | *Speciosa* | Hartmann 34813 (HBG) | South Africa: Western Cape-Uniondale | **LR030592** | **LR030782** | **LR030876** | **LR030684** | **LR030978** |

| **Species** | **Section** | **Voucher** | **Origin** | **ITS** | **rps16-trnK** | **trnQ-rps16** | **rpl16** | **trnS-G** |
|---|---|---|---|---|---|---|---|---|
| *Drosanthemum zygophylloides* (L.Bolus) L.Bolus | | Klak 830 (BOL) | South Africa: Western Cape-Piketberg | --- | --- | KF131863 | KF132156 | KF133123 |
| *Drosanthemum zygophylloides* (L.Bolus) L.Bolus | | Mucina 130216-4 | South Africa: Western Cape-Piketberg | LR030991 | LR031150 | LR031102 | LR031055 | LR031210 |
| **Outgroup** | **Clade in Klak et al. (2013)** | | | | | | | |
| *Antegibbaeum fissoides* (Haw.) Schwantes ex H.Wulff | L3 | Klak 308 (BOL) | South Africa: Western Cape | AJ438227 | -- | KF131867 | KF132159 | KF133127 |
| *Antimima ventricosa* (L.Bolus) H.E.K.Hartmann | L2 | Klak 475 (BOL) | South Africa: Northern Cape | AJ438225 | -- | JN896434 | KF132163 | JN896381 |
| *Braunsia geminata* (Haw.) L.Bolus | L2 | Klak 205 (BOL) | South Africa | AJ438228 | -- | KF131875 | KF132168 | KF133135 |
| *Carpobrotus muirii* (L.Bolus) L.Bolus | L1 | Klak 706 (BOL) | South Africa: Western Cape | AJ438230 | -- | KF131877 | KF132170 | KF133137 |
| *Cephalophyllum inaequale* L.Bolus | L1 | Hartmann 7883 (HBG) | South Africa: Western Cape | HG324002 | HG323950 | HG323976 | LR031056 | LR031211 |
| *Chasmatophyllum musculinum* (Haw.) Dinter & Schwantes | L3 | Bolduan s.n. | ex hort. ZSS (994202/0) | HG324003 | HG324003 | HG323977 | LR031057 | LR031212 |
| *Cheiridopsis pearsonii* N.E.Br. | I | Bruyns 9504 (BOL) | South Africa | -- | -- | KF131885 | KF132178 | KF133145 |
| *Cheiridopsis rostrata* (L.) N.E.Br. | I | Powell 88 (NBG) | South Africa: Western Cape | -- | -- | KY635278 | KY635037 | KY635358 |
| *Conophytum calculus* (A.Berger) N.E.Br. | I | Klak 1641 (BOL) | South Africa: Western Cape | -- | -- | KF131887 | KF132180 | KF133147 |
| | | Ritz s.n. | ex hort. | FN386499 | -- | -- | -- | -- |
| *Corpuscularia lehmannii* (Eckl. & Zeyh.) Schwantes | F | Klak 353 (BOL) | South Africa | -- | -- | KF131889 | KF132182 | KF133149 |
| *Corpuscularia lehmannii* (Eckl. & Zeyh.) Schwantes | | Hartmann & Kremling 33693 (HBG) | South Africa: Eastern Cape | AJ582942 | -- | -- | -- | -- |
| *Deilanthe peersii* (L.Bolus) N.E.Br. | L4 | Klak 1694 (BOL) | South Africa: Western Cape | -- | -- | KF131891 | KF132184 | KF133151 |
| *Deilanthe peersii* (L.Bolus) N.E.Br. | | Hartmann & Dehn 15486 (HBG) | South Africa: Eastern Cape | LR030992 | LR031151 | -- | -- | -- |

| **Species** | **Section** | **Voucher** | **Origin** | **ITS** | **rps16-trnK** | **trnQ-rps16** | **rpl16** | **trnS-G** |
|---|---|---|---|---|---|---|---|---|
| *Delosperma echinatum* (Lam.) Schwantes | F | Hartmann & Kremling 33682 (HBG) | South Africa: Eastern Cape | HG324005 | HG323953 | HG323979 | **LR031067** | **LR031225** |
| *Delosperma esterhuyseniae* L.Bolus | F | Bruyns 7141 (BOL) | South Africa | AJ438213 | -- | JN896438 | KF132187 | JN896385 |
| *Dicrocaulon brevifolium* N.E.Br. | B | Hartmann 1480 (HBG) | South Africa: Western Cape | LR030993 | LR031152 | LR031103 | LR031058 | LR031213 |
| *Dicrocaulon microstigma* (L.Bolus) Ihlenf. | B | Ihlenfeldt & Hartmann 4529 (HBG) | South Africa: Western Cape | HG324007 | HG323955 | HG323981 | LR031059 | LR031214 |
| *Disphyma dunsdonii* L.Bolus | C | Klak 808 (BOL) | South Africa | -- | -- | KF131902 | KF132194 | KF133162 |
| *Disphyma dunsdonii* L.Bolus | | Hartmann & Bayer 34630 (HBG) | South Africa: Western Cape | LR030994 | LR031153 | -- | -- | -- |
| *Drosanthemopsis diversifolia* (L.Bolus) Klak | H | Hartmann et al. 26169 (HBG) | South Africa: Western Cape | HG324013 | HG323961 | HG323987 | LR031060 | LR031215 |
| *Drosanthemopsis diversifolia* (L.Bolus) Klak | H | Klak 1743 (BOL) | South Africa | -- | -- | KF131856 | KF132149 | KF133116 |
| *Drosanthemopsis vaginata* (L.Bolus) Rauschert | H | Wisura 882 (BOL) | South Africa | -- | -- | KF131923 | KF132215 | KF133184 |
| *"Drosanthemum" pulverulentum* (Haw.) Schwantes | L1 | Klak 1847 (BOL) | South Africa: Western Cape | -- | -- | KF131860 | KF132152 | KF133120 |
| *Erepsia inclaudens* (Haw.) Schwantes | L1 | Bruyns 6847 (BOL) | South Africa: Western Cape | AJ438234 | -- | KF131907 | KF132200 | KF133168 |
| *Enarganthe octonaria* (L.Bolus) N.E.Br. | I | Klak 491 (BOL) | South Africa: Northern Cape | -- | -- | JN896433 | KF132198 | JN896380 |
| *Enarganthe octonaria* (L.Bolus) N.E.Br. | | Hartmann 7505 (HBG) | South Africa: Northern Cape | LR030995 | LR031154 | -- | -- | -- |
| *Esterhuysenia mucronata* (L.Bolus) Klak | L2 | Klak 709 (BOL) | South Africa: Western Cape | AJ438255 | | KF131908 | KF132201 | KF133169 |
| *Gibbaeum hortenseae* (N.E.Br.) Thiede & Klak | E | Klak 1979 (BOL) | South Africa | -- | -- | KF131913 | KF132205 | KF133174 |
| *Gibbaeum pachypodium* (Kensit) L.Bolus | E | Klak 380 (BOL) | South Africa | -- | -- | KF131914 | KF132206 | KF133175 |
| *Glottiphyllum cruciatum* (Haw.) N.E.Br. | D | Bruyns 8207 | South Africa | -- | -- | KF131915 | KF132207 | KF133176 |

| Species | Section | Voucher | Origin | ITS | rps16-trnK | trnQ-rps16 | rpl16 | trnS-G |
|---|---|---|---|---|---|---|---|---|
| *Glottiphyllum cruciatum* (Haw.) N.E.Br. | | Hartmann 8809 (HBG) | South Africa: Western Cape | LR030996 | LR031155 | -- | -- | -- |
| *Hartmanthus pergamentaceus* (L.Bolus) S.A.Hammer | K | Klak 243 (BOL) | Namibia | -- | -- | KF131919 | KF132211 | KF133180 |
| *Hartmanthus pergamentaceus* (L.Bolus) S.A.Hammer | K | Klak 1992 (BOL) | | -- | -- | KF131920 | KF132212 | KF133181 |
| *Ihlenfeldtia excavata* (L.Bolus) H.E.K.Hartmann | I | Wisura 1926 (BOL) | South Africa | -- | -- | KF131921 | KF132214 | KF133182 |
| *Ihlenfeldtia excavata* (L.Bolus) H.E.K.Hartmann | | Hartmann et al. 20587 (HBG) | South Africa: Northern Cape | LR030997 | LR031156 | -- | -- | -- |
| *Jacobsenia hallii* L.Bolus | J | Ihlenfeldt & Hartmann 4463 (HBG) | South Africa: Northern Cape | HG324024 | HG323973 | HG323999 | LR031061 | LR031216 |
| *Jacobsenia kolbei* (L.Bolus) L.Bolus & Schwantes | J | Klak 1819 (BOL) | South Africa | -- | -- | KF131922 | -- | KF133183 |
| *Jacobsenia kolbei* (L.Bolus) L.Bolus & Schwantes | | Ihlenfeldt & Hartmann 5060 (HBG) | South Africa: Western Cape | AJ582951 | -- | -- | -- | -- |
| *Lampranthus bicolor* (L.) N.E.Br. | L1 | Klak 543 (BOL) | South Africa: Western Cape | AJ438250 | -- | JN896437 | KF132221 | JN896384 |
| *Lithops julii* (Dinter & Schwantes) N.E.Br. ssp. fulleri (N.E.Br.) Fearn | L1 | Klak 701 (BOL) | South Africa: Northern Cape | AJ438218 | -- | KF131934 | KF132227 | KF133195 |
| *Malephora lutea* (Haw) Schwantes | D | Klak 664 (BOL) | South Africa: Western Cape | -- | -- | KF131936 | KF132229 | KF133197 |
| *Meyerophytum globosum* (L.Bolus) Ihlenf. | G | Hammer sub HBG 6314 (BOL) | South Africa | -- | -- | KF131939 | KF132087 | KF133200 |
| *Meyerophytum meyeri* (Schwantes) Schwantes | G | Rawe s.n. (BOL) | South Africa | -- | -- | KF131940 | KF132233 | KF133201 |
| *Meyerophytum meyeri* (Schwantes) Schwantes | | Ihlenfeldt & Hartmann 5046 (HBG) | South Africa: Northern Cape | LR030998 | LR031157 | -- | -- | -- |
| *Mitrophyllum clivorum* (N.E.Br.) Schwantes | G | Klak 1987 (BOL) | South Africa | -- | -- | KF131941 | KF132234 | KF133202 |

| Species | Section | Voucher | Origin | ITS | rps16-trnK | trnQ-rps16 | rpl16 | trnS-G |
|---|---|---|---|---|---|---|---|---|
| *Mitrophyllum clivorum* (N.E.Br.) Schwantes | | Ihlenfeldt & Hartmann 4681 (HBG) | South Africa: Northern Cape | LR030999 | LR031158 | -- | -- | -- |
| *Monilaria moniliformis* (Thunb.) Ihlenf. & Jürgens | B | Klak 787 (BOL) | South Africa: Western Cape | -- | -- | KF131942 | KF132235 | KF133203 |
| *Monilaria moniliformis* (Thunb.) Ihlenf. & Jürgens | | Ihlenfeldt & Hartmann 4213 (HBG) | South Africa: Western Cape | LR031000 | LR031159 | -- | -- | -- |
| *Nananthus aloides* (Haw.) Schwantes | L4 | Klak 1797 (BOL) | South Africa: North West Province | -- | -- | KF131946 | KF132238 | KF133207 |
| *Nananthus aloides* (Haw.) Schwantes | | Hartmann 32029 (HBG) | South Africa: Northern Cape | LR031001 | LR031160 | -- | -- | -- |
| *Odontophorus marlothii* N.E.Br. | I | Hartmann 7582 (HBG) | South Africa: Northern Cape | HG324025 | HG323974 | HG324000 | LR031062 | LR031217 |
| *Odontophorus marlothii* N.E.Br. | I | Klak 862 (BOL) | South Africa: Northern Cape | -- | -- | KF131951 | KF132243 | KF133212 |
| *Oscularia deltoides* (L.) Schwantes | L1 | Klak 215 (BOL) | South Africa | AJ438215 | -- | KF131955 | KF132246 | KF133216 |
| *Pleiospilos simulans* (Marloth) N.E.Br. | L4 | Bruyns 4988 (BOL) | South Africa: Eastern Cape | -- | -- | KF131959 | KF132250 | KF133220 |
| *Polymita steenbokensis* H.E.K.Hartmann | I | Bruyns 8267 (BOL) | South Africa: Northern Cape | -- | -- | KF131960 | KF132251 | KF133221 |
| *Polymita steenbokensis* H.E.K.Hartmann | | Hartmann & Bayer 78032 (HBG) | South Africa: Northern Cape | LR031002 | LR031161 | -- | -- | -- |
| *Prepodesma orpenii* (N.E.Br.) N.E.Br. | L4 | Klak 1800 (BOL) | South Africa: North West Province | -- | -- | KF131961 | -- | KF133222 |
| *Prepodesma orpenii* (N.E.Br.) N.E.Br. | | Hartmann & Dehn 19666 (HBG) | South Africa: Northern Cape | LR031003 | LR031162 | -- | -- | -- |
| *Roosia grahambeckii* (Van Jaarsv.) Van Jaarsv. | -- | VanJaarsveld s.n. (NBG) | South Africa: Western Cape | LR031004 | LR031163 | LR031104 | LR031063 | LR031218 |
| *Ruschia maxima* (Haw.) L.Bolus | L1 | Klak 704 (BOL) | South Africa | AJ438222 | -- | KF131969 | KF132258 | KF133231 |
| *Schlechteranthus hallii* L.Bolus | I | Klak 259 (BOL) | South Africa: Northern Cape | -- | -- | KF131980 | KF132268 | KF133242 |

| Species | Section | Voucher | Origin | ITS | rps16-trnK | trnQ-rps16 | rpl16 | trnS-G |
|---|---|---|---|---|---|---|---|---|
| *Schlechteranthus hallii* L.Bolus | | Hartmann 8276 (HBG) | South Africa: Northern Cape | LR031005 | -- | -- | -- | -- |
| *Scopelogena bruynsii* Klak | L1 | Klak 462 (BOL) | South Africa: Western Cape | AJ438258 | -- | KF131982 | KF132269 | KF133244 |
| *Smicrostigma viride* (Haw.) N.E.Br. | L2 | Klak 180 (BOL) | South Africa: Western Cape | AJ438259 | -- | JN896432 | KF132270 | JN896379 |
| *Vlokia ater* S.A.Hammer | L3 | Hammer & Vlok 1181 (BOL) | South Africa: Western Cape | AJ438260 | -- | KF131992 | KF132279 | KF133254 |
| *Wooleya farinosa* (L.Bolus) L.Bolus | L1 | Hartmann 7213 (HBG) | South Africa: Northern Cape | HG324026 | HG323975 | HG324001 | LR031064 | LR031219 |
| *Zeuktophyllum suppositum* (L.Bolus) N.E.Br. | L2 | Klak 375 (BOL) | South Africa: Western Cape | AJ438262 | -- | KF131994 | KF132281 | KF133256 |
| **Dorotheantheae** | | | | | | | | |
| *Cleretum bellidiforme* (Burm.f.) G.D.Rowley | | Klak 627 (BOL)/ Klak 1534 (BOL) | South Africa: South Western Cape | KF132145 | JN896383 | JN896436 | -- | JN896503 |
| *Cleretum papulosum* (L.f.) L.Bolus | | Klak 1487 (BOL) | South Africa: Northern Cape | KF132146 | JN896408 | JN896461 | -- | JN896507 |
| *Cleretum rourkei* L. Bolus | | Klak 1524 (BOL) | South Africa: Northern Cape | -- | JN896500 | JN896454 | -- | JN896500 |
| **Apatesieae** | | | | | | | | |
| *Conicosia pugioniformis* (L.) N.E.Br. ssp. *muirii* (N.E.Br.) Ihlenf. & Gerbaulet | | Klak 1570 | South Africa: South Western Cape | KF132144 | JN896409 | JN896462 | -- | JN896508 |
| *Conicosia pugioniformis* (L.) N.E.Br. | | Juergens sn. | South Africa | HG324004 | HG323952 | HG323978 | -- | -- |
