## Supplementary material for "Phylogenetic relationships in the southern African genus *Drosanthemum* (Ruschioideae, Aizoaceae)": S2 Character recoding

Examples of character re-coding used for intra-clade haplotype analyses:

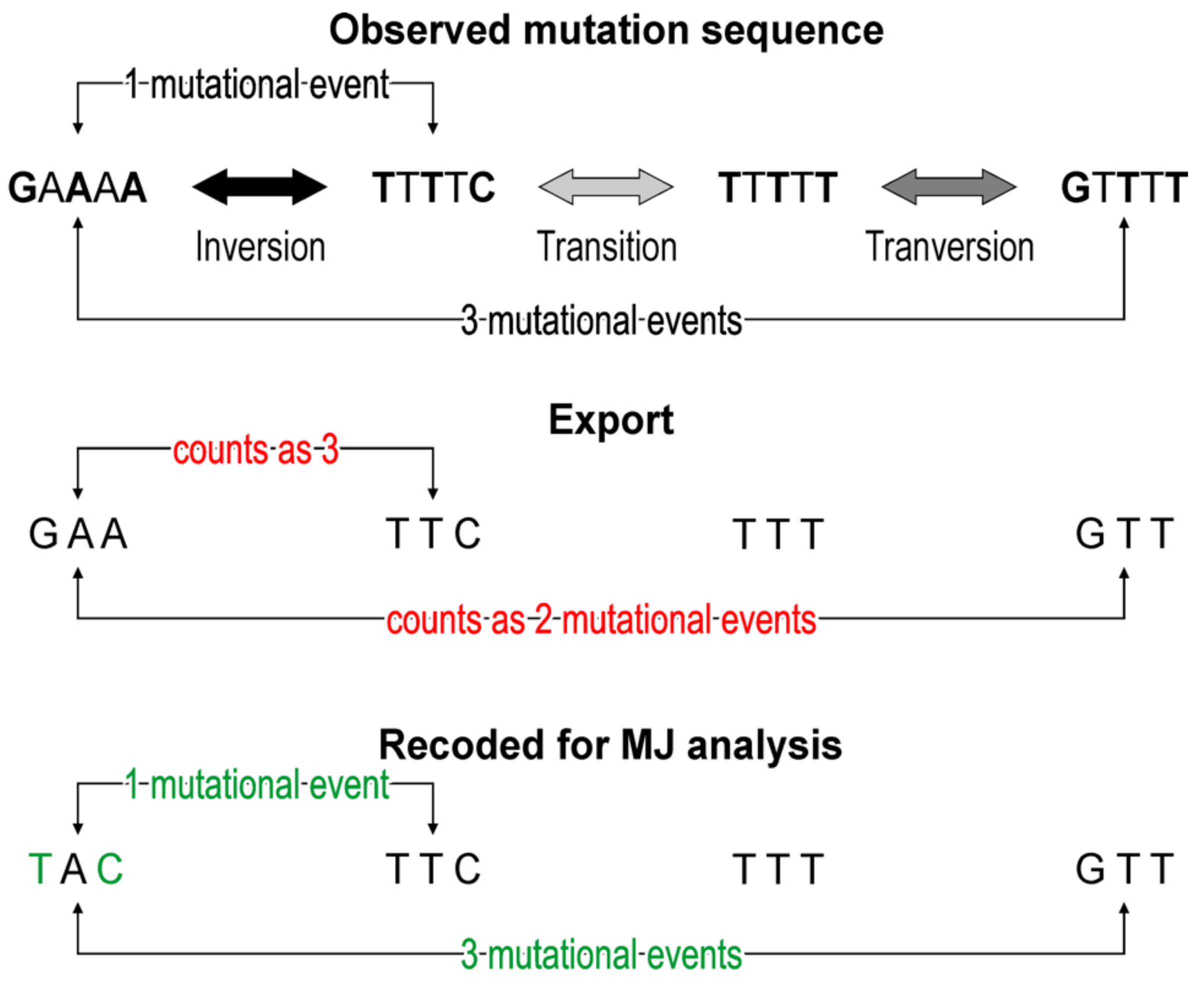


**Figure S2 (A)**: **Examples of character re-coding**. Pseudo-loop motif (inversion with secondary point mutations), as found in the complementary ('pseudo-hairpin') region of the *trnK-rps16* in Clade III and IV (characters 7–9).

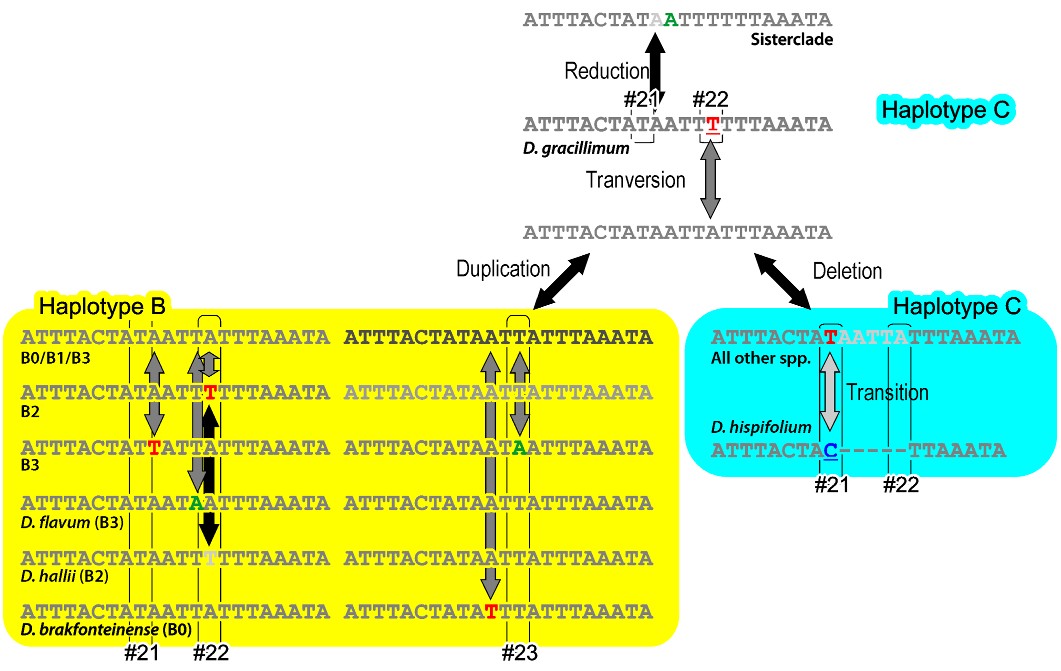


Oligonucleotide motif at 5' end of *trnS-trnG* in Clades 5 and 6 (chars # 21–23)

**Figure S2 (B)**: **Examples of character re-coding**. Oligonucleotide motif at 5' end of *trnS-trnG* in Clades V and VI (characters 21–23).

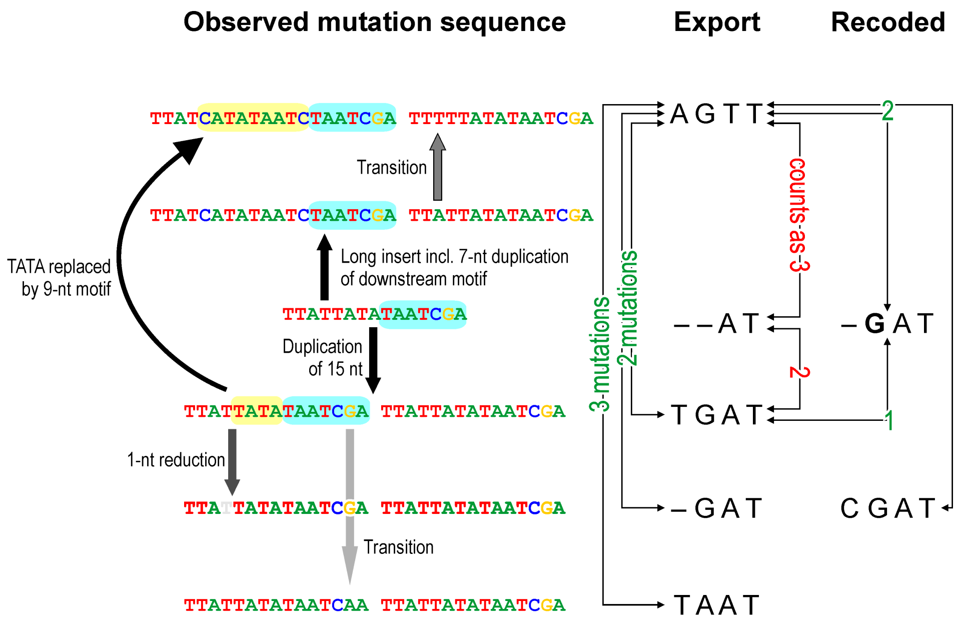


Indel pattern with secondary SNP in the centre of *trnS-trnG* in Clades 7 to 9 (chars # 13–16)

**Figure S2 (C)**: **Examples of character re-coding**. Indel pattern with secondary SNP in the centre of *trnS-trnG* in Clades VII–IX (characters 13–16).
