## Supplementary material for "Phylogenetic relationships in the southern African genus *Drosanthemum* (Ruschioideae, Aizoaceae)": S3 Statistical parsimony (SP) network analyses of ITS sequence variation

### Drosanthemum rDNA networks

Alastair Potts

16 Jul 2019

```
library(ape)
library(pegas)
library(dplyr)
library(ggplot2)
library(scales)
library(flextable)
```

### SP network for ITS variants of (cp) clade I to IX

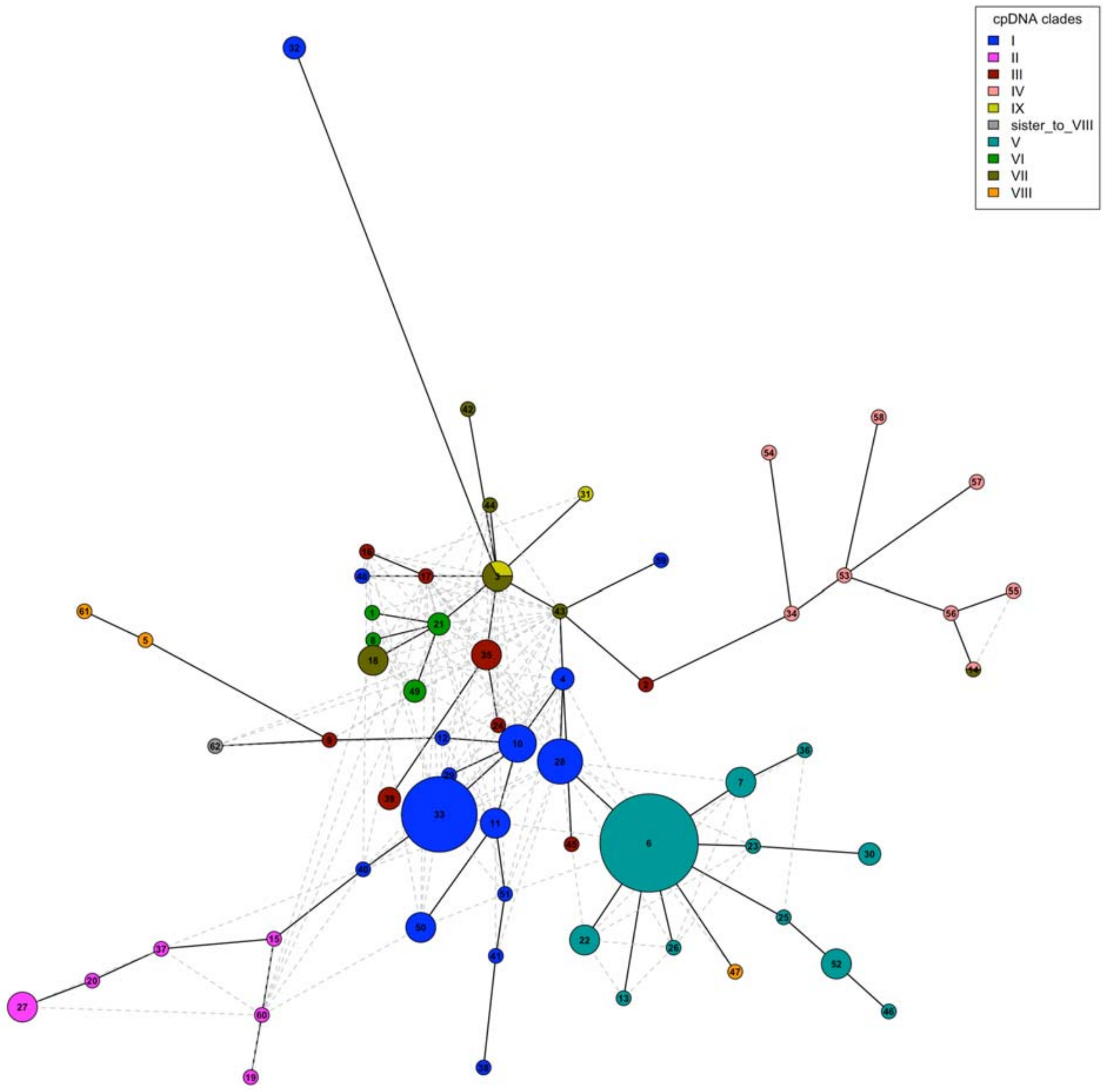


The star-shape is generally taken to be an indication of a bottleneck event with subsequent expansion (if we were looking at species phylogeography).

Given its central location, haplotype 3 might indicate the ancestral haplotype (rather than the common 6…)

| ITS_hap | Accession |
|---|---|
| 1 | Drosanthemum_acuminatum_HH_34608 |
| 2 | Drosanthemum_acutifolium_HH_32406 |
| 3 | Drosanthemum_anemophilum_VanJaarsveld_sn |
| 3 | Drosanthemum_calycinum_HH_32211 |
| 3 | Drosanthemum_papillatum_HH_34624 |
| 4 | Drosanthemum_archeri_HH_31774 |
| 4 | Drosanthemum_archeri_HH_31788 |
| 5 | Drosanthemum_asperulum_HH_34502 |

| ITS_hap | Accession |
|---|---|
| 6 | Drosanthemum_attenuatum_aff_HH_34650 |
| 6 | Drosanthemum_attenuatum_HH_11826 |
| 6 | Drosanthemum_austricola_HH_34695 |
| 6 | Drosanthemum_boerhavii_HH_34697 |
| 6 | Drosanthemum_cereale_HH_34491 |
| 6 | Drosanthemum_chrysum_HH_34631 |
| 6 | Drosanthemum_hallii_HH_34610 |
| 6 | Drosanthemum_insolitum_HH_34596 |
| 6 | Drosanthemum_pulchrum_HH_34712 |
| 6 | Drosanthemum_stokoei_Bayer_7454 |
| 6 | Drosanthemum_strictifolium_HH_34480 |
| 6 | Drosanthemum_uniondalensis_HH_34813 |
| 7 | Drosanthemum_aureopurpureum_HH_34478 |
| 7 | Drosanthemum_brakfonteinii_HH_34688 |
| 7 | Drosanthemum_micans_HH_34597 |
| 8 | Drosanthemum_bicolor_HH_32459 |
| 9 | Drosanthemum_brevifolium_HH_32242 |
| 10 | Drosanthemum_candens_HH_25014 |
| 10 | Drosanthemum_floribundum_HH_34403 |
| 10 | Drosanthemum_hispidum_HH_33866 |
| 10 | Drosanthemum_sp_Drosanthemum_HH_30568 |
| 11 | Drosanthemum_candens_HH_30541 |
| 11 | Drosanthemum_dejagerae_HH_31812 |
| 11 | Drosanthemum_sp_Drosanthemum_HH_33182 |
| 12 | Drosanthemum_candens_HH_31897 |
| 13 | Drosanthemum_cereale_HH_34492 |
| 14 | Drosanthemum_crassum_HH_25896 |
| 14 | Drosanthemum_crassum_HH_25896 |
| 15 | Drosanthemum_curtophyllum_HH_32673 |
| 16 | Drosanthemum_cymiferum_HH_25811 |
| 17 | Drosanthemum_cymiferum_HH_31686 |
| 18 | Drosanthemum_cymiferum_HH_32250 |
| 18 | Drosanthemum_expersum_HH_34590 |
| 18 | Drosanthemum_sp_Quastea_HH_34598 |
| 19 | Drosanthemum_delicatulum_HH_32462 |
| 20 | Drosanthemum_dipageae_HH_34409 |
| 21 | Drosanthemum_ecclesianum_HH_34814 |
| 21 | Drosanthemum_ecclesianum_HH_34815 |
| 22 | Drosanthemum_edwardsiae_HH_34648 |
| 22 | Drosanthemum_lavisii_HH_34693 |
| 22 | Drosanthemum_speciosum_HH_34619 |
| 23 | Drosanthemum_edwardsiae_HH_34651 |
| 24 | Drosanthemum_erigeriflorum_HH_26196 |
| 25 | Drosanthemum_flammeum_HH_34460 |
| 26 | Drosanthemum_flavum_HH_34706 |
| 27 | Drosanthemum_fourcadei_HH_33900 |
| 27 | Drosanthemum_nodosum_HH_34404 |
| 27 | Drosanthemum_sp_Xamera_HH_34586 |
| 28 | Drosanthemum_framesii_HH_32339 |

| ITS_hap | Accession |
|---|---|
| 28 | Drosanthemum_glabrescens_HH_32256 |
| 28 | Drosanthemum_marinum_HH_32205 |
| 28 | Drosanthemum_prostratum_aff_HH_34316 |
| 28 | Drosanthemum_prostratum_HH_34592 |
| 29 | Drosanthemum_hispidum_HH_34262 |
| 30 | Drosanthemum_hispifolium_HH_32206 |
| 30 | Drosanthemum_hispifolium_HH_34587 |
| 31 | Drosanthemum_inornatum_HH_32654 |
| 32 | Drosanthemum_intermedium_HH_32200 |
| 32 | Drosanthemum_intermedium_HH_32439 |
| 33 | Drosanthemum_latipetalum_HH_31568 |
| 33 | Drosanthemum_latipetalum_HH_31605 |
| 33 | Drosanthemum_latipetalum_HH_32217 |
| 33 | Drosanthemum_muirii_HH_30321 |
| 33 | Drosanthemum_nordenstamii_HH_31533 |
| 33 | Drosanthemum_oculatum_HH_31710 |
| 33 | Drosanthemum_schoenlandianum_HH_25751 |
| 33 | Drosanthemum_sp_Drosanthemum_HH_32218 |
| 33 | Drosanthemum_subplanum_HH_32259 |
| 34 | Drosanthemum_lique_HH_34447 |
| 35 | Drosanthemum_luederitzii_HH_25972 |
| 35 | Drosanthemum_luederitzii_HH_26095 |
| 35 | Drosanthemum_obibense_HH_25979 |
| 36 | Drosanthemum_micans_LeRoux_83_2 |
| 37 | Drosanthemum_montaguense_HH_31824 |
| 38 | Drosanthemum_muiri_HH_32209 |
| 39 | Drosanthemum_nollothense_HH_31547 |
| 39 | Drosanthemum_nollothense_HH_31569 |
| 40 | Drosanthemum_oculatum_HH_31680 |
| 41 | Drosanthemum_opacum_HH_32212 |
| 42 | Drosanthemum_papillatum_HH_34456 |
| 43 | Drosanthemum_papillatum_HH_34607 |
| 44 | Drosanthemum_papillatum_HH_34614 |
| 45 | Drosanthemum_parvifolium_HH_30399 |
| 46 | Drosanthemum_pustulatum_HH_34698 |
| 47 | Drosanthemum_quadratum_HH_34503 |
| 48 | Drosanthemum_schoenlandianum_Bruyns_7172 |
| 49 | Drosanthemum_semiglobosum_HH_34593 |
| 49 | Drosanthemum_thudichumii_HH_34714 |
| 50 | Drosanthemum_sp_Drosanthemum_HH_25365 |
| 50 | Drosanthemum_sp_Drosanthemum_HH_32392neu |
| 50 | Drosanthemum_sp_Drosanthemum_HH_34496 |
| 51 | Drosanthemum_sp_Drosanthemum_HH_34472 |
| 52 | Drosanthemum_sp_Ossicula_HH_34699 |
| 52 | Drosanthemum_striatum_HH_29013 |
| 52 | Drosanthemum_striatum_HH_34720 |
| 53 | Drosanthemum_sp_Vespertina_HH_26170a |
| 54 | Drosanthemum_sp_Vespertina_HH_31059 |
| 55 | Drosanthemum_sp_Vespertina_HH_32378 |

| ITS_hap | Accession |
|---|---|
| 56 | Drosanthemum_sp_Vespertina_HH_32404 |
| 57 | Drosanthemum_sp_Vespertina_HH_33490 |
| 58 | Drosanthemum_sp_Vespertina_HH_34804 |
| 59 | Drosanthemum_sp_Vespertina_Mucina_161005_14 |
| 60 | Drosanthemum_subclausum_HH_25737 |
| 61 | Drosanthemum_tetramerum_HH_34488 |
| 62 | Drosanthemum_zygophylloides_Mucina_130216_4 |

### Mismatch distribution

The histogram shows the observed distribution (frequencies) of pairwise distances from the set of ITS sequences. The lines show an empirical density estimate (in blue) and the expected distribution under stable population (Rogers and Harpending 1992).

```
MMD(as.DNAbin(nex))
```

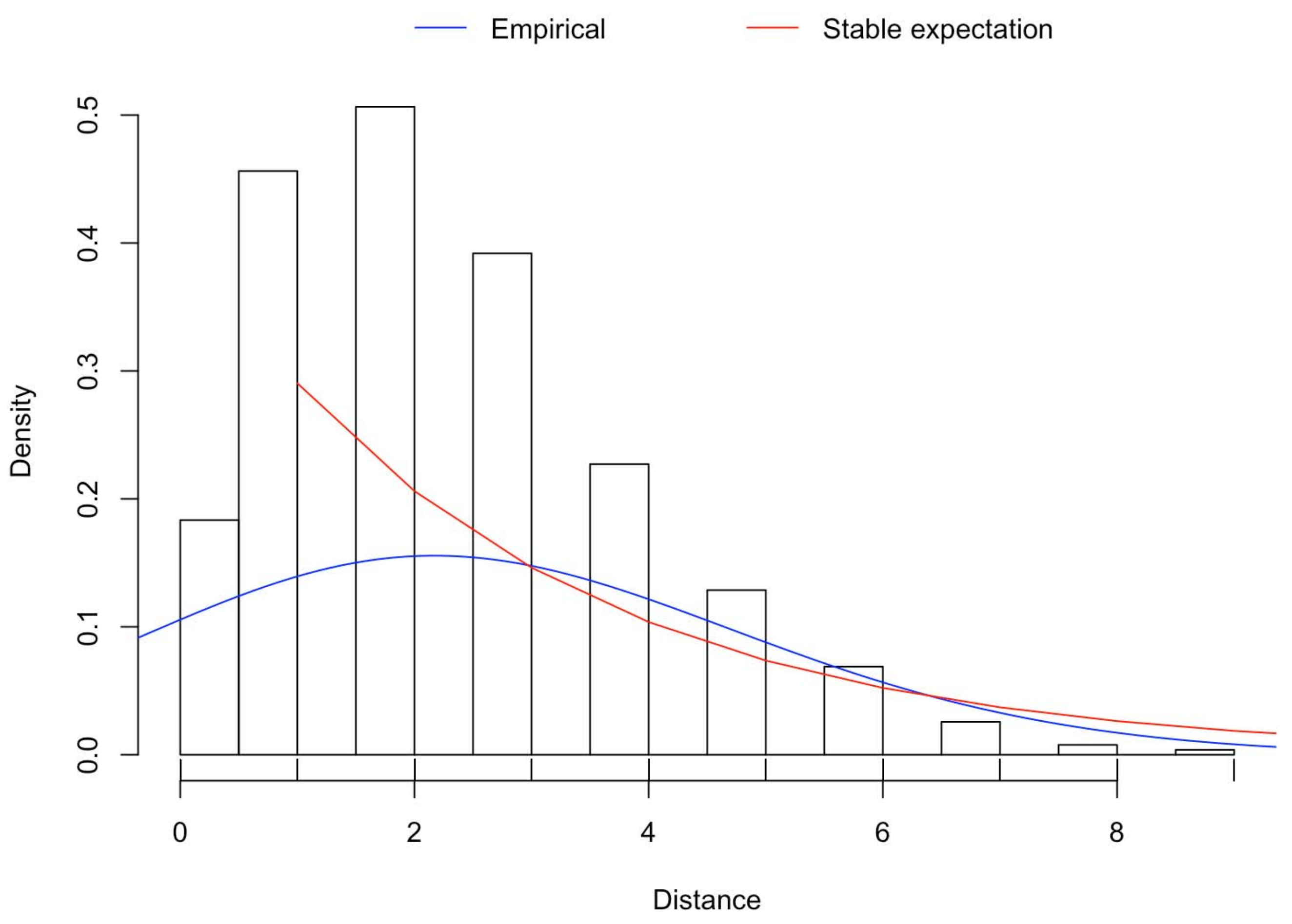


The mismatch distribution indicates an expanding “population”, in this case, probably radiations/emeregence of new species.

### Subclade networks

### SP network for ITS variants of clade I to IV, VIII, and IX

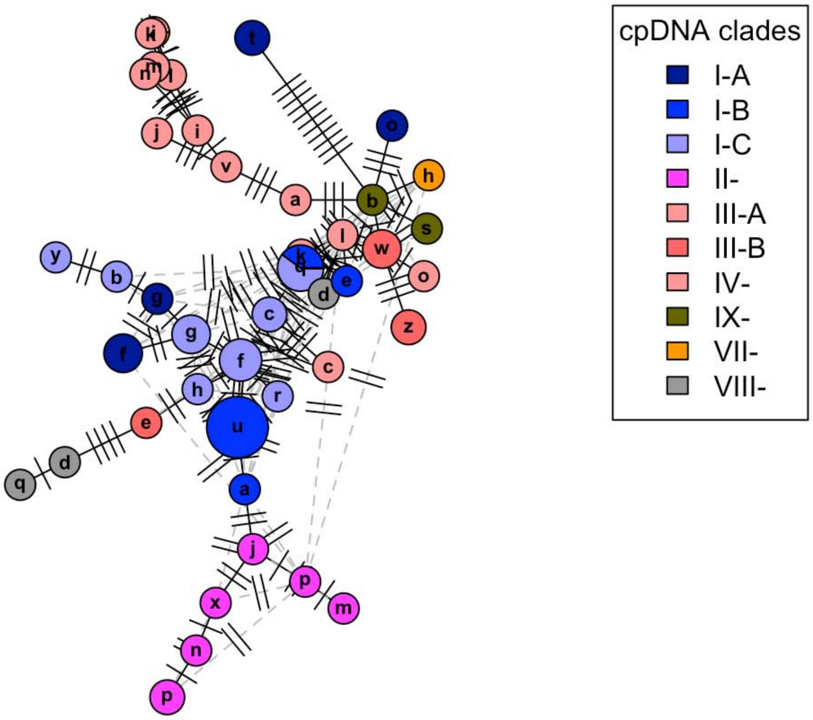

### SP network for ITS variants of clade I to IV

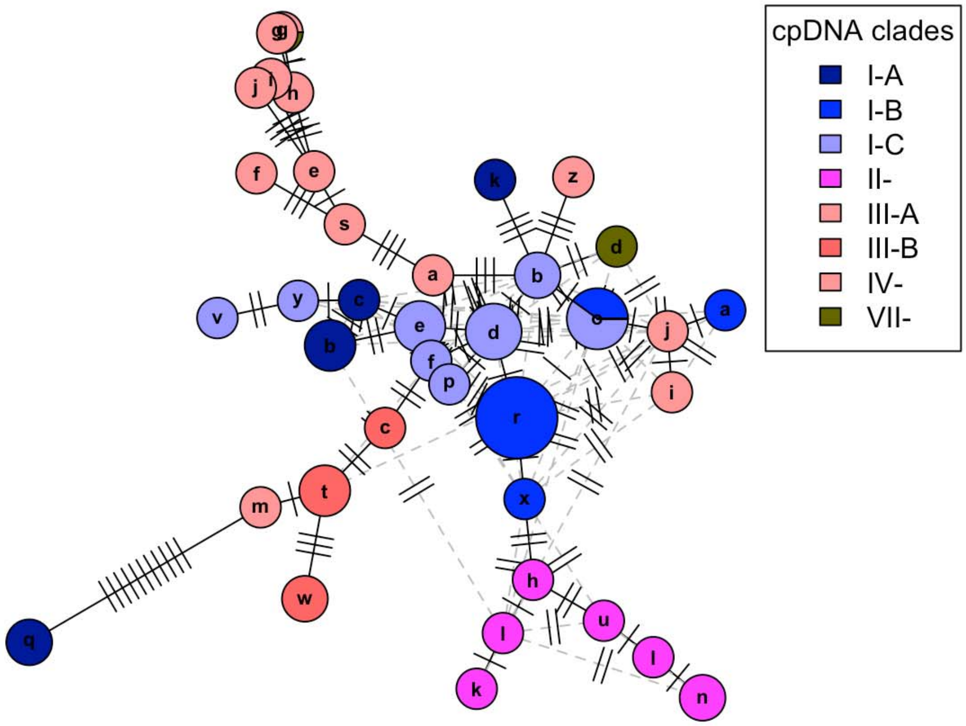

### SP network for ITS variants of clade I and II

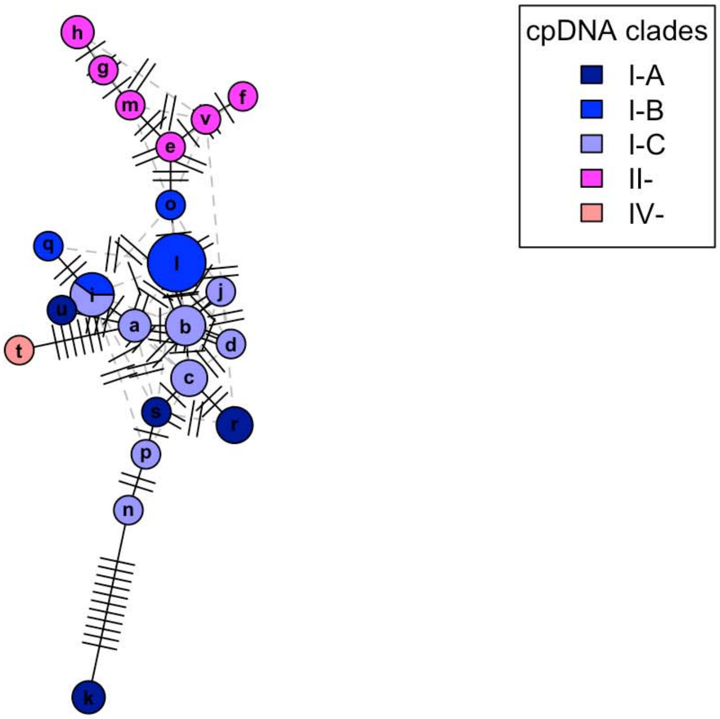

### SP network for ITS variants of clade III and IV

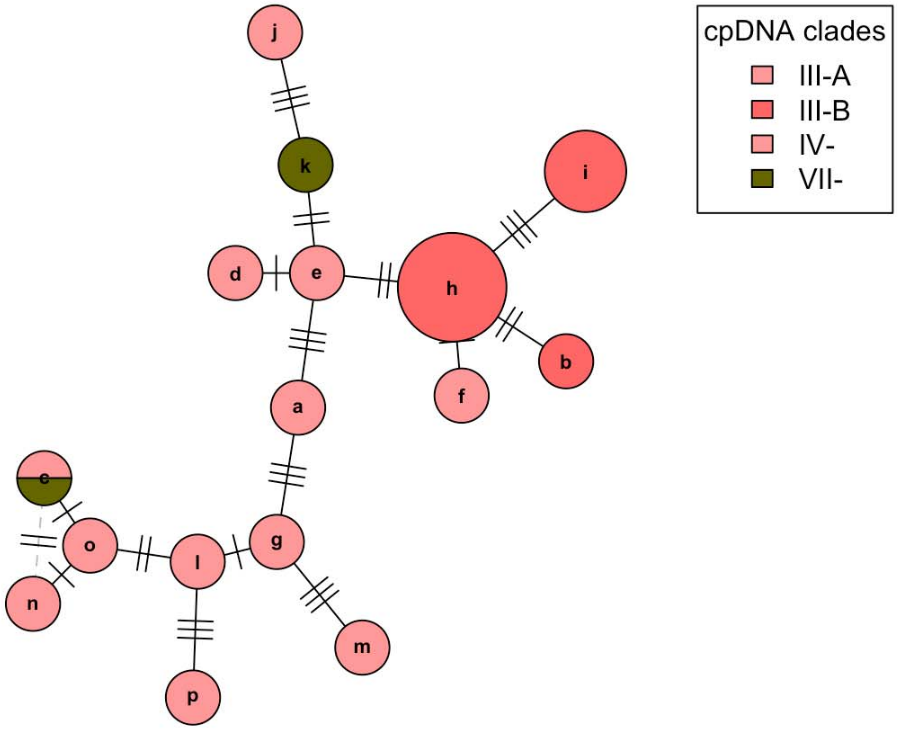

### SP network for ITS variants of clade V to VII

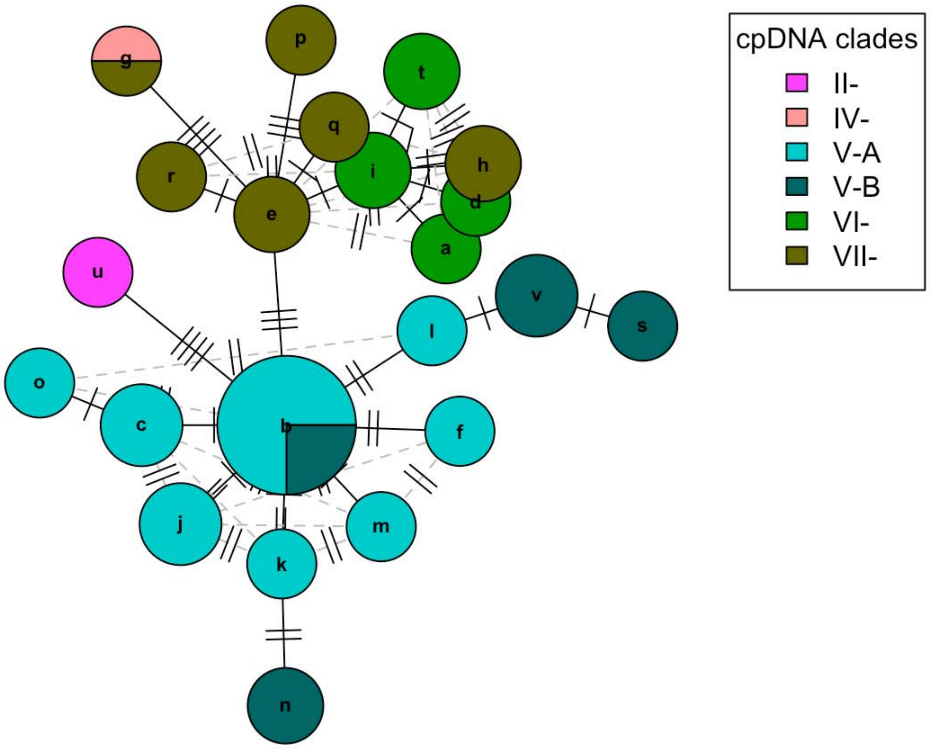

### SP network for ITS variants of clade VIII, sister to VIII, and IX

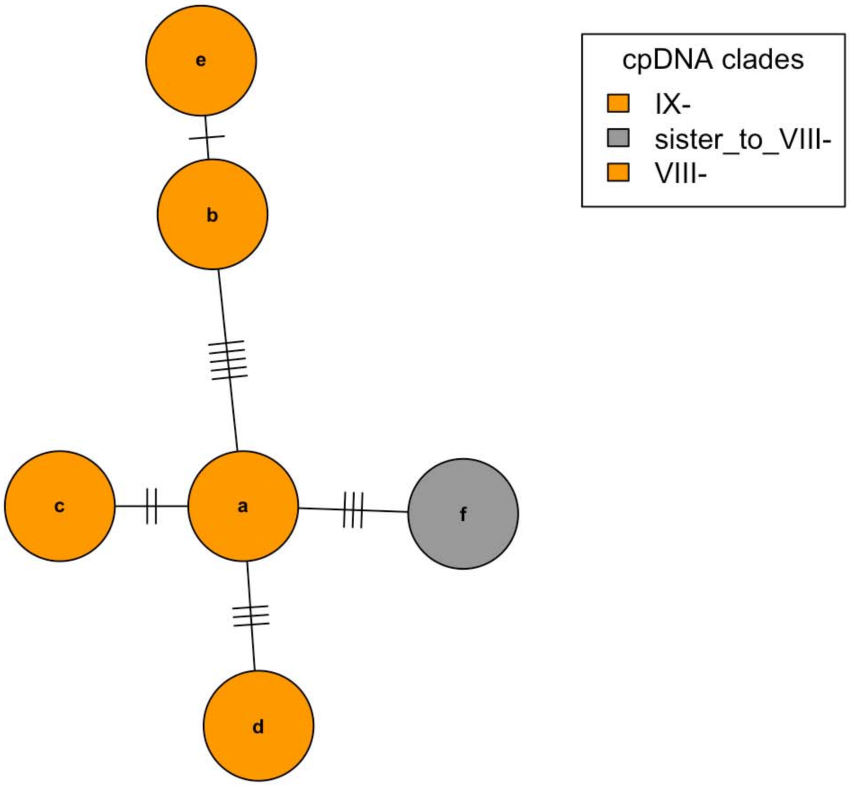
