## Supplementary material for "Phylogenetic relationships in the southern African genus *Drosanthemum* (Ruschioideae, Aizoaceae)": S4 Outgroup EPA

**Contains:**

1) Results of maximum likelihood analysis of the partitioned "Ruschioideae" cpDNA dataset (Fig. S4.1).
2) Results of outgroup-EPA analyses (Berger, Krompass & Stamatakis 2011) detailing the five potential root positions in *Drosanthemum* with queried Ruschieae outgroup taxa supporting a respective scenario (Figs. S4.2–6). Likelihood weight ratios for the queried outgroup taxa and probability estimates (Table S4). Raw data, code and full analysis output for all partitioning schemes and marker combinations are available at Dryad, Liede-Schumann et al. 2019.

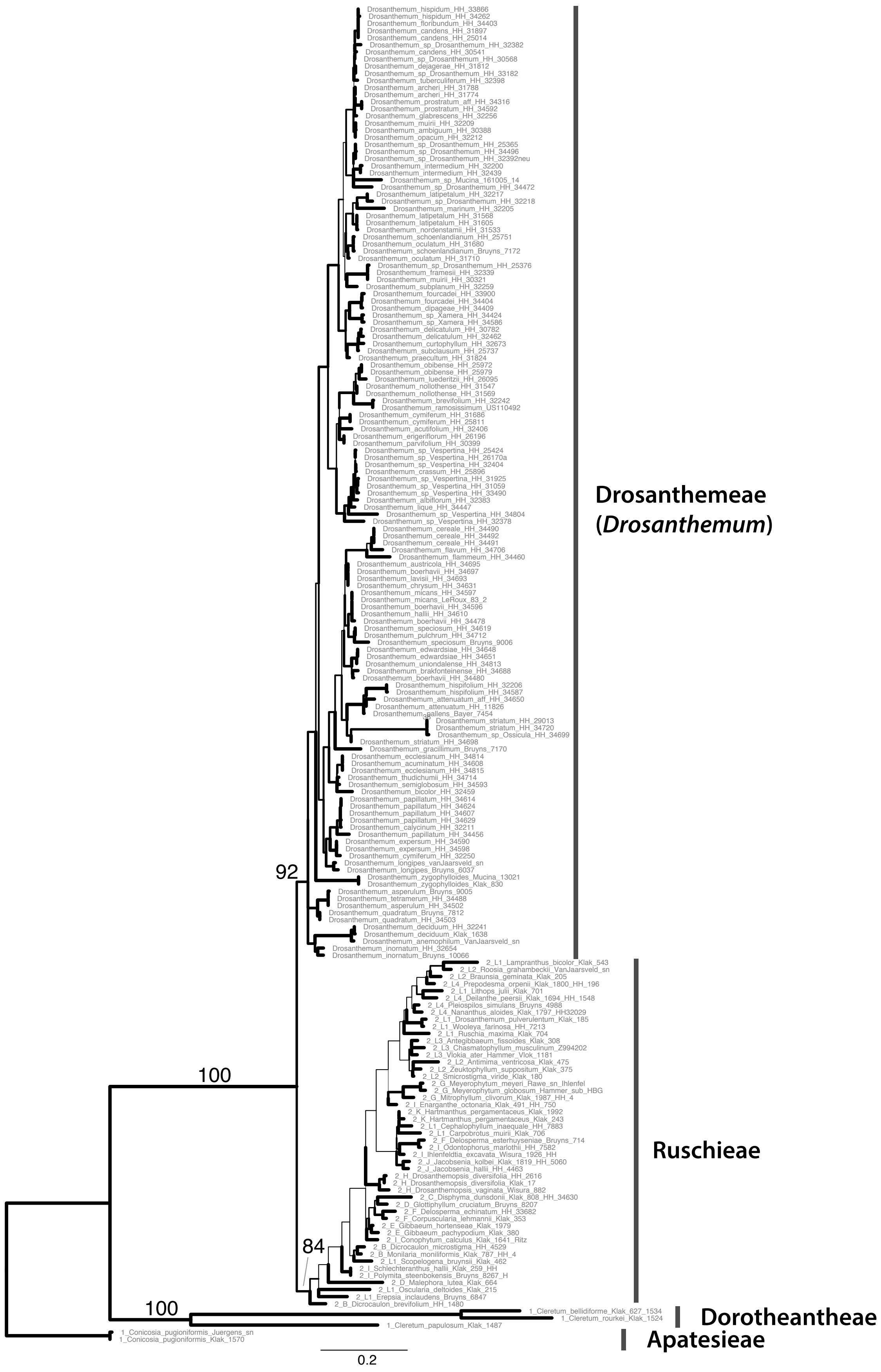


**Figure S4.1**: **Phylogeny Ruschioideae.** ML tree inferred by partitioned analysis of the cpDNA sequence data for Ruschioideae. The tribes of Ruschioideae are annotated.

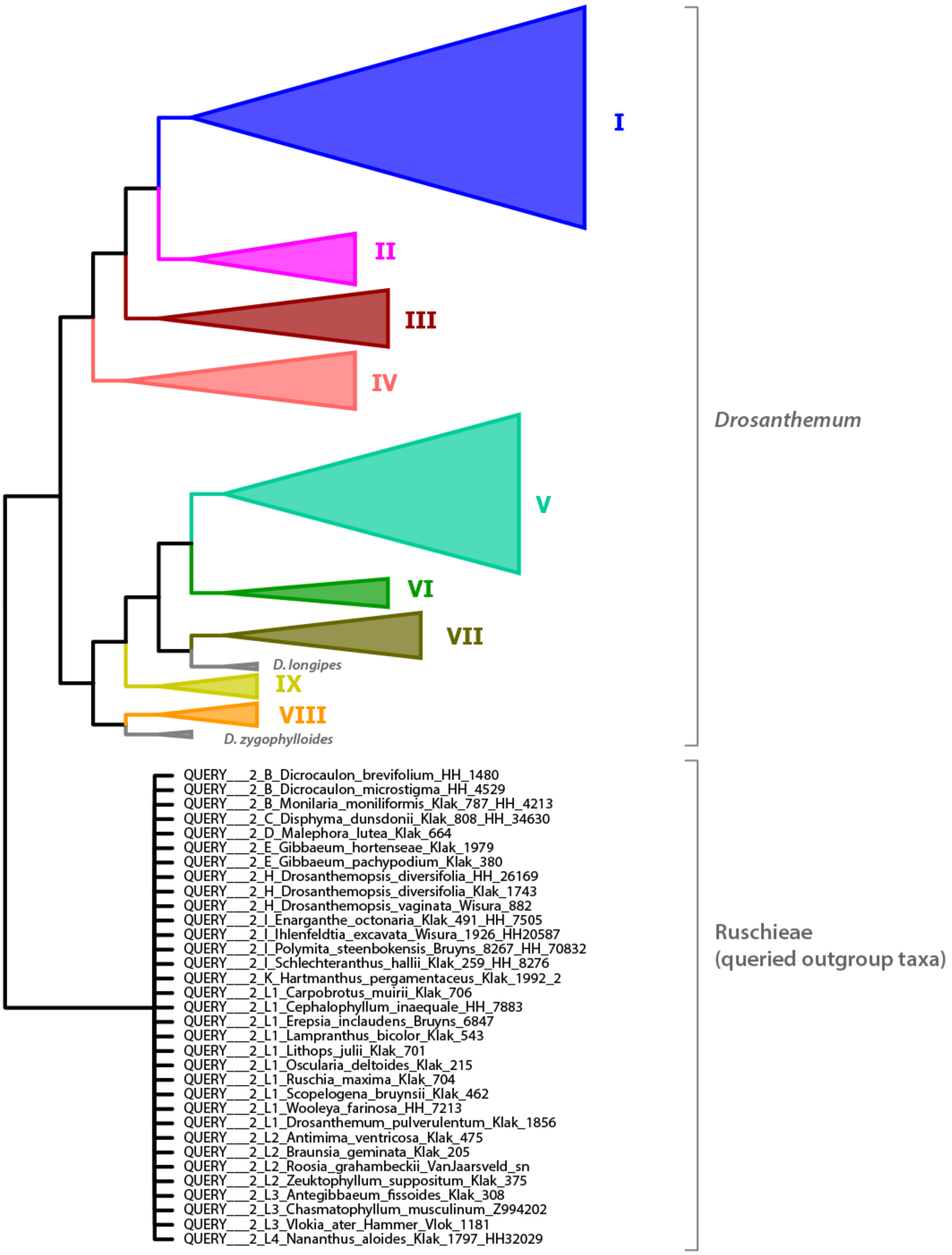


**Figure S4.2**: ***Drosanthemum* root position scenario 1**. Mean probability estimate 0.260.

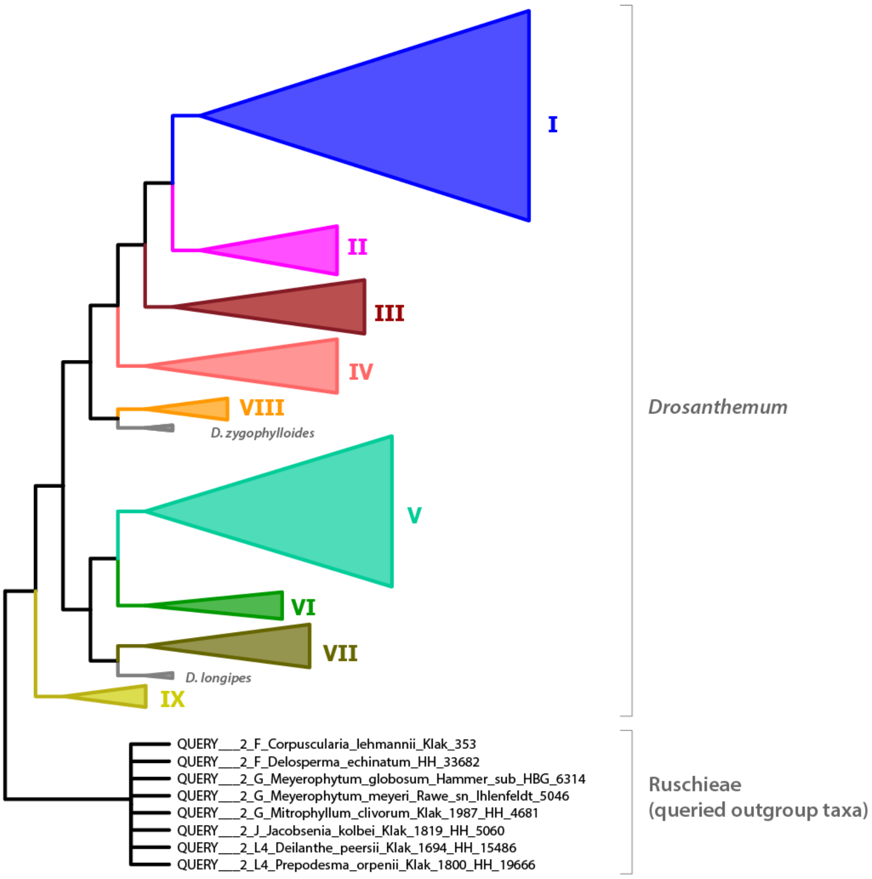


**Figure S4.3**: ***Drosanthemum* root position scenario 2**. Mean probability estimate 0.232.

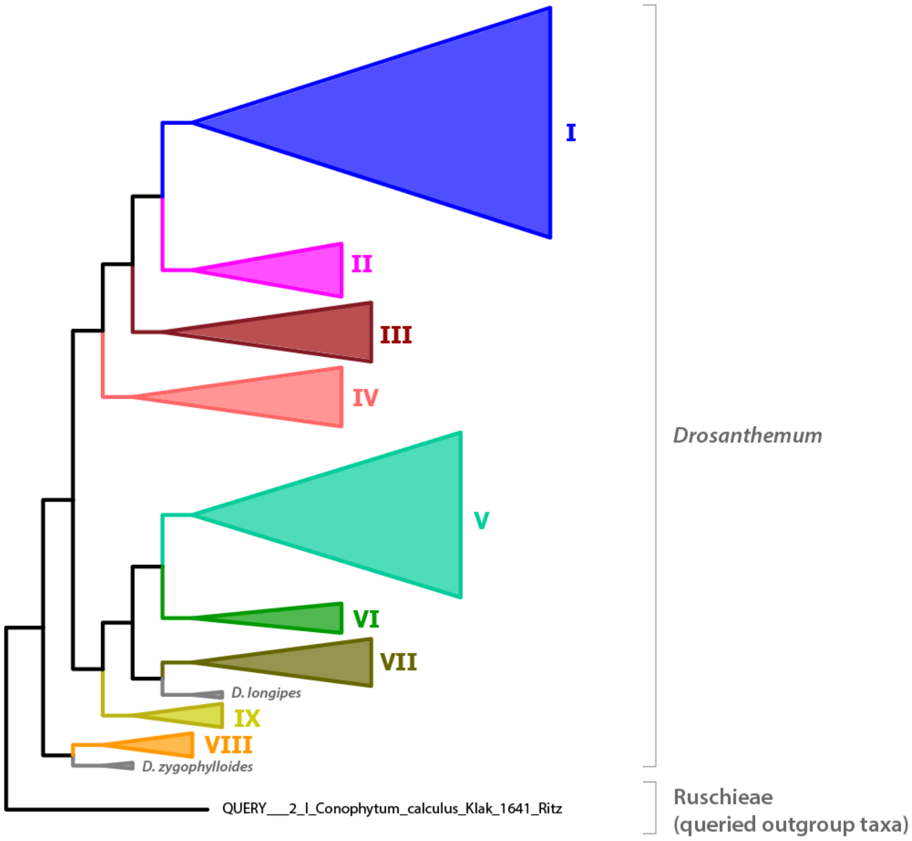


**Figure S4.4**: ***Drosanthemum* root position scenario 3**. Mean probability estimate 0.143.

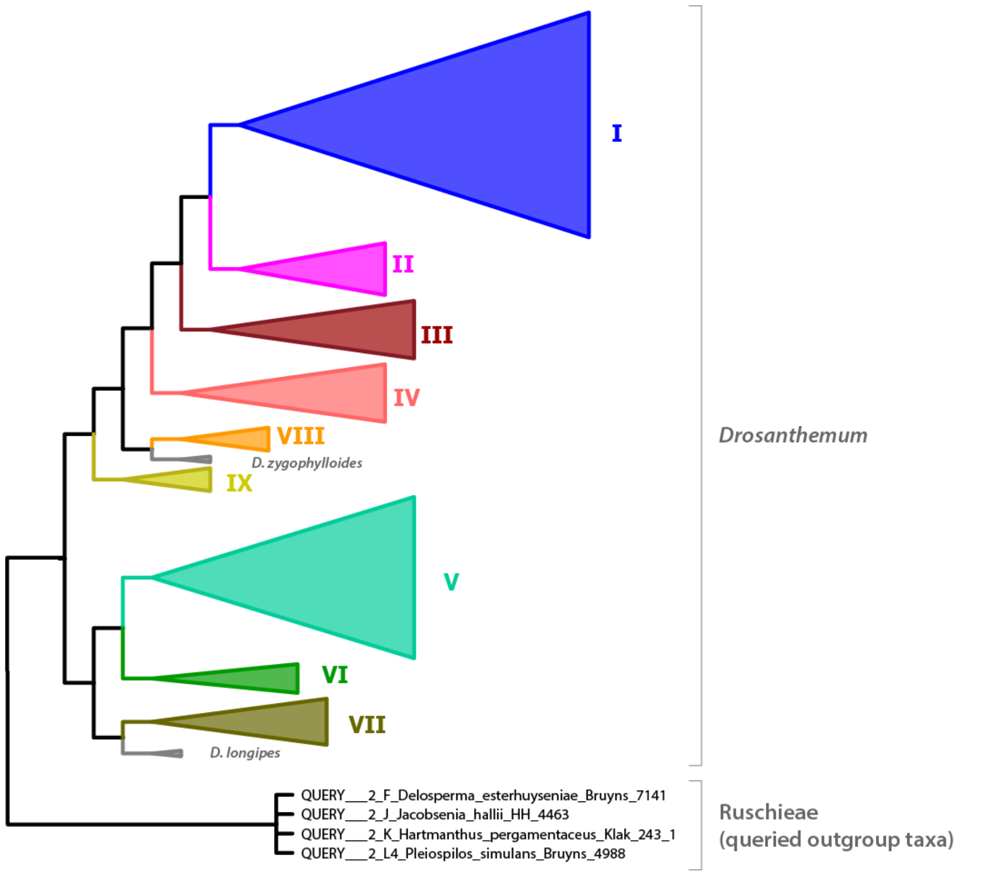


**Figure S4.5**: ***Drosanthemum* root position scenario 4**. Mean probability estimate 0.138.

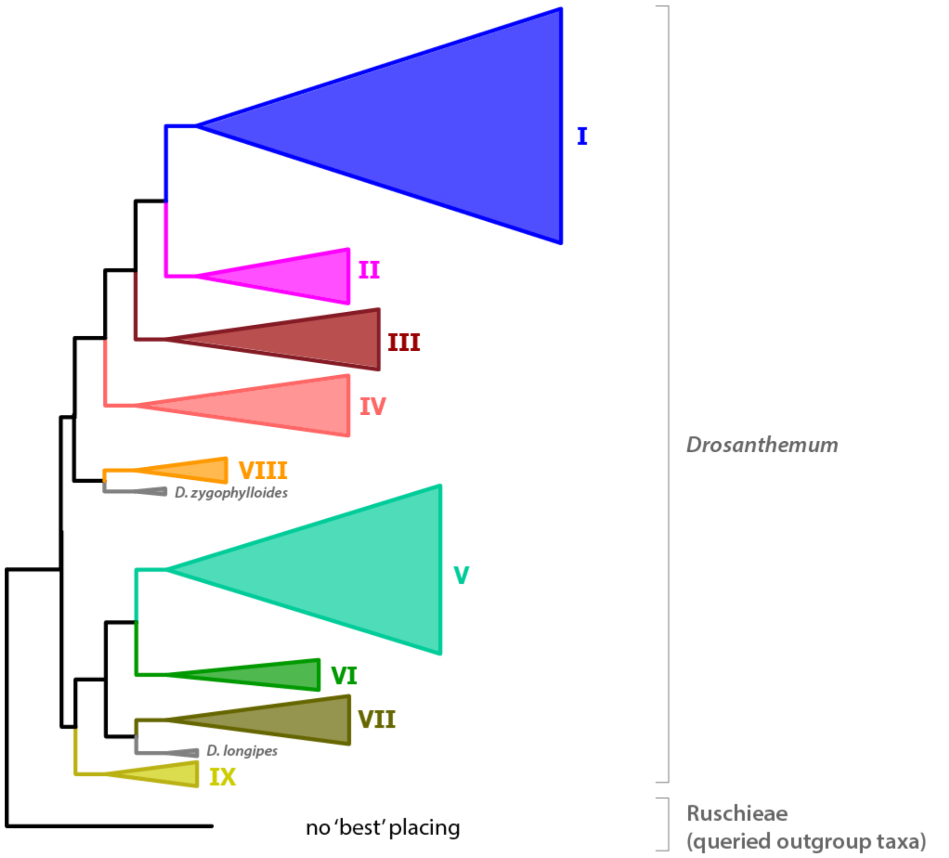


**Figure S4.6**: ***Drosanthemum* root position scenario 5**. Mean probability estimate 0.137.

**Table S4. Results of outgroup EPA**. *Drosanthemum* rooting scenarios S1–S11 detailing inferred root position by node numbers, number of queried outgroup taxa supporting a scenario, the mean probability estimates and the queried outgroup taxa by their likelihood weight ratio per scenario. Numbers in bold fond indicate the supporting taxa per scenario (i.e. highest likelihood weight ratio).

| **Query results** | **S 1** | **S 2** | **S 3** | **S 4** | **S 5** | **S 6** | **S 7** | **S 8** | **S 9** | **S 10** | **S 11** |
|---|---|---|---|---|---|---|---|---|---|---|---|
| Node Nº [a] | 180 | 182 | 232 | 186 | 181 | 233 | 234 | 223 | 222 | 187 | 224 |
| Internal node Nº [b] | I94 | I96 | I105 | I198 | I95 | I199 | I202 | I178 | I177 | I106 | I181 |
| # placings (supporting taxa) | 33 | 8 | 1 | 4 | 0 | 2 | 0 | 0 | 1 | 0 | 0 |
| Mean probability estimate | 0.2596 | 0.2321 | 0.1430 | 0.1377 | 0.1372 | 0.0297 | 0.0192 | 0.0165 | 0.0102 | 0.0016 | 0.0010 |
| Query taxa | | | | | | | | | | | |
| 2_B_Dicrocaulon_brevifolium_HH_1480 | **0.1789** | 0.1777 | 0.1775 | 0.1781 | 0.1777 | 0.0162 | | | | 0.0599 | |
| 2_B_Dicrocaulon_microstigma_HH_4529 | **0.1932** | 0.1915 | 0.1915 | 0.1917 | 0.1916 | 0.0175 | 0.0158 | | | | |
| 2_B_Monilaria_moniliformis_Klak_787_HH_4213 | **0.1910** | 0.1895 | 0.1894 | 0.1898 | 0.1896 | 0.0175 | 0.0160 | | | | |
| 2_C_Disphyma_dunsdonii_Klak_808_HH_34630 | **0.7077** | 0.0655 | 0.0655 | 0.0655 | 0.0655 | 0.0190 | | | | | |
| 2_D_Glottiphyllum_cruciatum_Bruyns_8207_HH_8809 | 0.0167 | | | 0.0170 | 0.0164 | | | 0.3408 | **0.5002** | 0.0182 | 0.0471 |
| 2_D_Malephora_lutea_Klak_664 | **0.1831** | 0.1816 | 0.1816 | 0.1817 | 0.1817 | 0.0195 | | 0.0416 | | | |
| 2_E_Gibbaeum_hortenseae_Klak_1979 | **0.1927** | 0.1907 | 0.1908 | 0.1909 | 0.1908 | 0.0197 | 0.0157 | | | | |
| 2_E_Gibbaeum_pachypodium_Klak_380 | **0.1923** | 0.1899 | 0.1899 | 0.1904 | 0.1899 | 0.0174 | 0.0157 | | | | |
| 2_F_Corpuscularia_lehmannii_Klak_353 | 0.0751 | **0.6587** | 0.0746 | 0.0747 | 0.0746 | 0.0224 | 0.0083 | | | | |
| 2_F_Delosperma_echinatum_HH_33682 | 0.0627 | **0.7306** | 0.0622 | 0.0624 | 0.0623 | | | | | | |
| 2_F_Delosperma_esterhuyseniae_Bruyns_7141 | 0.1900 | 0.1898 | 0.1899 | **0.1900** | 0.1899 | 0.0211 | 0.0175 | | | | |
| 2_G_Meyerophytum_globosum_Hammer_sub_HBG_6314 | 0.1046 | **0.5569** | 0.1041 | 0.1042 | 0.1042 | 0.0100 | 0.0093 | | | | |
| 2_G_Meyerophytum_meyeri_Rawe_sn_Ihlenfeldt_5046 | 0.0951 | **0.5368** | 0.0942 | 0.0943 | 0.0943 | 0.0720 | 0.0083 | | | | |
| 2_G_Mitrophyllum_clivorum_Klak_1987_HH_4681 | 0.0949 | **0.5406** | 0.0952 | 0.0945 | 0.0944 | 0.0654 | 0.0089 | | | | |
| 2_H_Drosanthemopsis_diversifolia_HH_26169 | **0.1811** | 0.1794 | 0.1794 | 0.1795 | 0.1794 | 0.0731 | 0.0169 | | | | |
| 2_H_Drosanthemopsis_diversifolia_Klak_1743 | **0.1913** | 0.1894 | 0.1894 | 0.1896 | 0.1895 | 0.0190 | 0.0180 | | | | |
| 2_H_Drosanthemopsis_vaginata_Wisura_882 | **0.1911** | 0.1893 | 0.1893 | 0.1894 | 0.1893 | 0.0188 | 0.0182 | | | | |
| 2_I_Conophytum_calculus_Klak_1641_Ritz | 0.1902 | 0.1898 | **0.1906** | 0.1899 | 0.1898 | 0.0208 | 0.0167 | | | | |
| 2_I_Enarganthe_octonaria_Klak_491_HH_7505 | **0.8948** | 0.0250 | 0.0250 | 0.0250 | 0.0250 | | | | | | |
| 2_I_Ihlenfeldtia_excavata_Wisura_1926_HH20587 | **0.8523** | 0.0349 | 0.0349 | 0.0349 | 0.0349 | | | | | | |
| 2_I_Odontophorus_marlothii_HH_7582 | 0.0464 | | 0.2988 | | | **0.3494** | 0.2969 | | | | |
| 2_I_Polymita_steenbokensis_Bruyns_8267_HH_70832 | **0.8817** | 0.0278 | 0.0278 | 0.0279 | 0.0279 | | | | | | |

| Query results | S 1 | S 2 | S 3 | S 4 | S 5 | S 6 | S 7 | S 8 | S 9 | S 10 | S 11 |
|---|---|---|---|---|---|---|---|---|---|---|---|
| 2_I_Schlechteranthus_hallii_Klak_259_HH_8276 | **0.1964** | 0.1857 | 0.1857 | 0.1859 | 0.1858 | 0.0284 | 0.0184 | | | | |
| 2_J_Jacobsenia_hallii_HH_4463 | 0.1894 | 0.1878 | 0.1879 | **0.2011** | 0.1879 | 0.0211 | 0.0192 | | | | |
| 2_J_Jacobsenia_kolbei_Klak_1819_HH_5060 | 0.0475 | **0.7941** | 0.0470 | 0.0471 | 0.0471 | 0.0100 | | | | | |
| 2_K_Hartmanthus_pergamentaceus_Klak_1992_2 | **0.1925** | 0.1908 | 0.1908 | 0.1909 | 0.1909 | 0.0206 | 0.0155 | | | | |
| 2_K_Hartmanthus_pergamentaceus_Klak_243_1 | 0.1876 | 0.1871 | 0.1871 | **0.1919** | 0.1871 | 0.0236 | 0.0210 | | | | |
| 2_L1_Carpobrotus_muirii_Klak_706 | **0.1922** | 0.1905 | 0.1905 | 0.1907 | 0.1906 | 0.0212 | 0.0158 | | | | |
| 2_L1_Cephalophyllum_inaequale_HH_7883 | **0.1799** | 0.1785 | 0.1785 | 0.1787 | 0.1786 | 0.0796 | 0.0162 | | | | |
| 2_L1_Drosanthemum_pulverulentum_Klak_1856 | **0.1903** | 0.1899 | 0.1899 | 0.1900 | 0.1900 | 0.0198 | 0.0175 | | | | |
| 2_L1_Erepsia_inclaudens_Bruyns_6847 | **0.1902** | 0.1886 | 0.1886 | 0.1888 | 0.1887 | 0.0201 | 0.0189 | | | | |
| 2_L1_Lampranthus_bicolor_Klak_543 | **0.4983** | 0.0251 | 0.0251 | 0.0251 | 0.0251 | | | 0.3913 | | | |
| 2_L1_Lithops_julii_Klak_701 | **0.1922** | 0.1904 | 0.1904 | 0.1906 | 0.1905 | 0.0196 | 0.0163 | | | | |
| 2_L1_Oscularia_deltoides_Klak_215 | **0.1916** | 0.1896 | 0.1896 | 0.1897 | 0.1897 | 0.0186 | 0.0178 | | | | |
| 2_L1_Ruschia_maxima_Klak_704 | **0.1842** | 0.1826 | 0.1826 | 0.1828 | 0.1827 | 0.0175 | | 0.0371 | | | |
| 2_L1_Scopelogena_bruynsii_Klak_462 | **0.1907** | 0.1899 | 0.1899 | 0.1901 | 0.1900 | 0.0207 | 0.0170 | | | | |
| 2_L1_Wooleya_farinosa_HH_7213 | **0.1931** | 0.1916 | 0.1916 | 0.1917 | 0.1916 | 0.0174 | 0.0163 | | | | |
| 2_L2_Antimima_ventricosa_Klak_475 | **0.1922** | 0.1903 | 0.1903 | 0.1905 | 0.1904 | 0.0193 | 0.0165 | | | | |
| 2_L2_Braunsia_geminata_Klak_205 | **0.8531** | 0.0344 | 0.0344 | 0.0345 | 0.0345 | | | | | | |
| 2_L2_Roosia_grahambeckii_VanJaarsveld_sn | **0.7860** | 0.0506 | 0.0507 | 0.0507 | 0.0507 | | | | | | |
| 2_L2_Smicrostigma_viride_Klak_180 | 0.1356 | 0.1344 | 0.1346 | 0.1346 | 0.1345 | **0.1882** | 0.1318 | | | | |
| 2_L2_Zeuktophyllum_suppositum_Klak_375 | **0.7438** | 0.0597 | 0.0597 | 0.0598 | 0.0598 | | | | | | |
| 2_L3_Antegibbaeum_fissoides_Klak_308 | **0.1921** | 0.1904 | 0.1904 | 0.1905 | 0.1905 | 0.0188 | 0.0167 | | | | |
| 2_L3_Chasmatophyllum_musculinum_Z994202 | **0.1911** | 0.1907 | 0.1907 | 0.1909 | 0.1908 | 0.0181 | 0.0182 | | | | |
| 2_L3_Vlokia_ater_Hammer_Vlok_1181 | **0.1919** | 0.1900 | 0.1900 | 0.1902 | 0.1901 | 0.0214 | 0.0168 | | | | |
| 2_L4_Deilanthe_peersii_Klak_1694_HH_15486 | 0.0108 | **0.9546** | 0.0107 | 0.0107 | 0.0107 | | | | | | |
| 2_L4_Nananthus_aloides_Klak_1797_HH32029 | **0.1921** | 0.1903 | 0.1903 | 0.1904 | 0.1903 | 0.0180 | 0.0184 | | | | |
| 2_L4_Pleiospilos_simulans_Bruyns_4988 | 0.1901 | 0.1900 | 0.1900 | **0.1908** | 0.1900 | 0.0211 | 0.0175 | | | | |
| 2_L4_Prepodesma_orpenii_Klak_1800_HH_19666 | 0.1188 | **0.5005** | 0.1177 | 0.1178 | 0.1177 | 0.0115 | 0.0107 | | | | |

a, Node No refers to the ML cpDNA tree (Fig 3), see Dryad Liede-Schumann et al. 2019, file d_RAxML_bestTree.6_cp.q.tre (can be viewed in e.g. R using ape::nodelabels).
b, Internal node No refers to the RAxML EPA output file detailing position of queried outgroup taxa, see Dryad Liede-Schumann et al. 2019, file RAxML_labelledTree.OGPA.q (can be viewed in e.g. Dendroscope).
